# Mechanochemical Feedback between Cell Shape and Intracellular Mechanics Revealed by a Finite-Element Framework

**DOI:** 10.64898/2026.07.03.736361

**Authors:** Alessandro Contri, Emmet A. Francis, André Massing, Padmini Rangamani

## Abstract

Cell shape and mechanics are intricately connected and tightly regulated by mechanochemical events including biochemical signaling, cytoskeletal remodeling, and plasma membrane mechanics. While experimental advances in microscopy have shed light on the intricate coordination involved in cell shape change in response to different cues, the ability to conduct three-dimensional simulations in realistic geometries remains an open computational challenge. In this work, we develop a finite-element framework that incorporates advection-diffusion-reaction equations coupled with equations governing the kinematics of a deformable interface representing the cell membrane. We applied this framework to three distinct coupled mechanochemical systems, each governed by geometric partial differential equations, resulting in large deformations of the interface. In all three examples, our simulations revealed the emergence of feedback between cellular signaling, cytoskeletal organization, and cell shape. In our first two sets of simulations, we observed that cell migration and neutrophil protrusion were regulated by membrane tension-mediated feedback. In our final application, we predicted shape changes of a dendritic spine starting from a realistic geometry, and found that the complex shape of the spine gives rise to localized regimes of actin cytoskeleton remodeling not previously observed with idealized geometries. Thus, our finite-element framework allows us to generate new mechanistic insights for biophysical problems.

---

Cells change their shape in response to environmental cues during development, wound healing, and differentiation [1–3]. These shape changes are driven by the integration of chemical and mechanical signals received at the plasma membrane, leading to changes in biochemical signaling within the cytosol and resulting in cytoskeletal reorganization [4–6] (Figure 1A). Remodeling of the cytoskeleton then leads to force generation at the plasma membrane, which itself is a deformable fluidic interface with resistance to bending [7–9] (Figure 1A). The literature is rich with experimental observations of such phenomena, including cell migration [10–16], subcellular shape changes that arise during different modes of membrane trafficking, including endocytosis [17, 18] and phagocytosis [19–22], droplet wetting [23–31], and even synaptic plasticity [32–34].

**Fig. 1:**
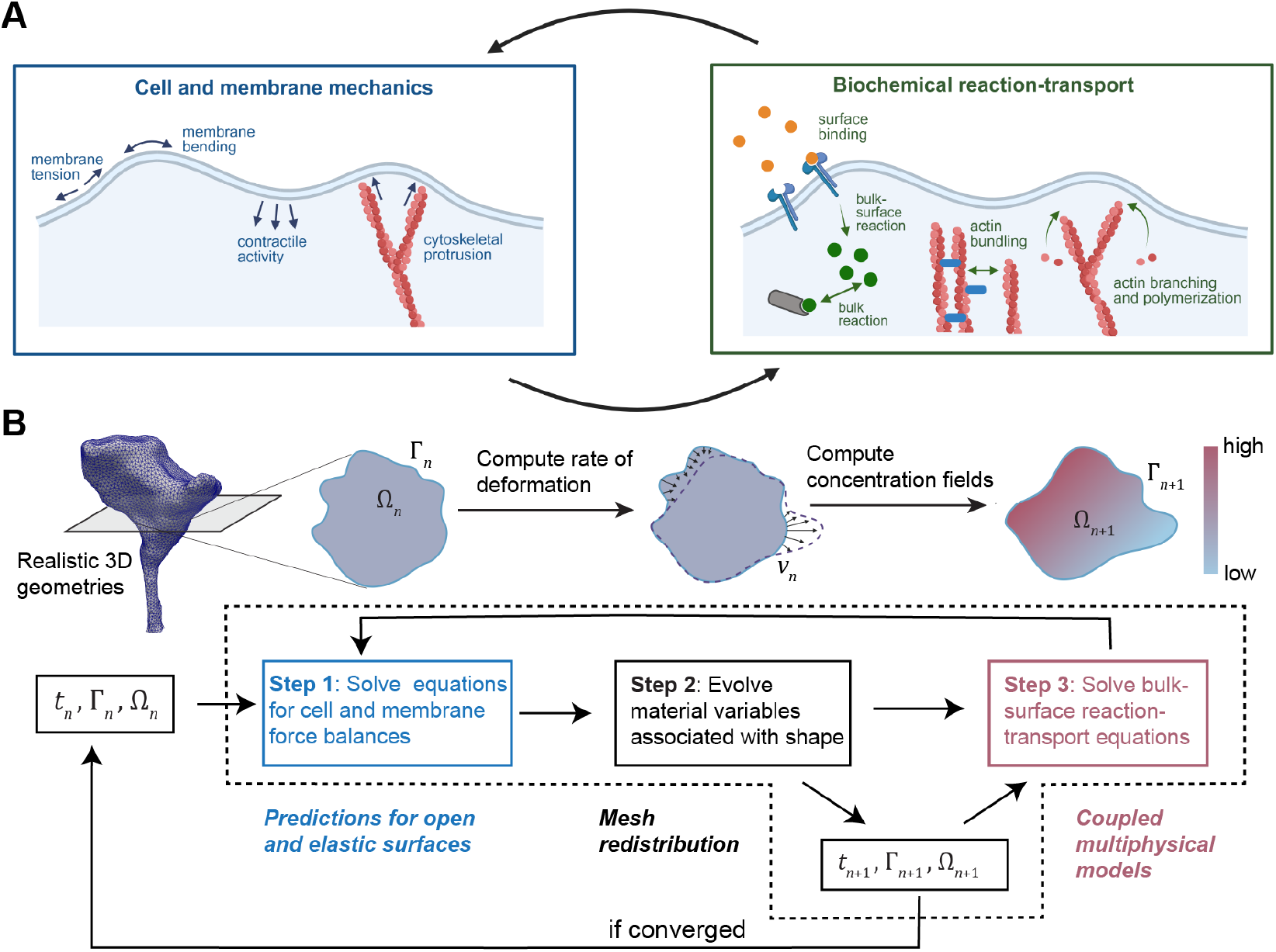
A continuum modeling framework for mechanochemical coupling in cells. A) The interplay between cytoskeletal and membrane mechanics results in changes to cell shape. These mechanical forces are directly informed by biochemical events including signal transduction and cytoskeletal remodeling. Resulting changes to the cell shape further influence the reactions establishing feedback between changes to cell shape and the underlying mechanochemical events. Schematic created with Biorender.com. B) A simulation framework for solving governing equations in cell biophysics. Step 1: Stable force-balance equations on surfaces dictate the physical shape evolution. Step 2: ALE update preserves mesh quality during geometrical shape evolution. Step 3: Structure-preserving reaction-transport equations on updated geometry provide interpretable results that feed back into the force balance.

Continuum modeling offers a rich landscape for studying such biophysical phenomena [7, 8, 35, 36] and generating new insights into the underlying biological mechanisms. These systems are modeled using systems of partial differential equations (PDEs) that couple advection-diffusion-reaction equations to the deformation of a free surface at the boundary. Additional complexity can arise from mixed-dimensional equations coupled by Robin boundary conditions that capture reactions at the interface of volumetric compartments and membranes [36, 37] and from the complex geometries of cells [36, 38]. The deformation of the cell membrane is often modeled using the Helfrich energy [39]. Many such models are considered in 2D to generate mechanistic insights and represent cells that have flattened during migration over flat substrates [12, 40, 41]. While this assumption is valid in certain cases, cells in 3D environments adopt much more complex geometries [38, 42, 43]. Understanding how realistic cell shapes in 3D can undergo large shape changes remains an open problem in biophysics. Furthermore, the development of robust numerical frameworks in a unified and stable setting to solve systems of PDEs describing mechanochemically coupled systems and to generate physically relevant simulations of cell shape changes remains an open challenge in the field of computational science.

One approach to solving these equations is to use Finite Element Methods (FEM). FEM frameworks have the advantage of a strong theoretical foundation and broad applicability to a wide range of mechanical problems [44–46]. However, schemes for generalized elastic membrane flows of the Helfrich type on surfaces with boundary that are provably robust and can handle large deformations are scarce in the literature. As exceptions, the schemes in [47, 48] are among the only ones to our knowledge that are able to handle both large deformations and surfaces with boundary in a full 3D setting. Coupled FEM frameworks for multiphysics problems on deformable membranes have also been explored in the literature [49–51], yielding important insights into the associated biological processes. There have been a number of recent advances in surface finite element method (SFEM) modeling [52] and its application to biophysical problems [49–51, 53, 54]. We note that the aforementioned studies often overlook convergence and stability properties in favor of computational efficiency. Moreover, the interpretability of results may be compromised when simulations fail to satisfy basic biophysical constraints such as conservation of mass and non-negativity of species concentrations. Finally, we note that none of these algorithms has been adapted to simulate PDEs in realistic cell shapes, which may pose unique meshing and computational challenges of their own.

In this work, we leverage the advances in FEM described above and build on our previous work [55] to develop a stable, finite-element-based framework that can be used to solve geometric PDEs with membrane reshaping on realistic meshes derived from experimental observations in the large deformation regime (Figure 1B). The key contributions of this work are as follows. We extend the algorithm for Helfrich flow presented in [48] by considering a heterogeneous spontaneous curvature that coevolves with the mean curvature equation. This extension maintains the energy-decaying property of the scheme and is an important step in working with realistic geometries. The resulting two-stage algorithm is then seamlessly integrated into our previous framework [55]. The key properties of the previous framework — stability, convergence, tunability, and interpretability — are preserved in the new framework. The result is a *unified finite element framework* applicable to realistic geometries undergoing large deformations where a true energetic ground state is guaranteed. We demonstrate the generalizability and applicability of this framework using three specific biophysical examples: cell migration, neutrophil chemotactic protrusion, and structural plasticity of synapses. In all these cases, our simulations reveal that the feedback between signaling, cytoskeletal remodeling, and cell shape change is governed by mechanochemical events that occur at the interface.

## 1 Results

### 1.1 A novel bulk-surface FEM framework to simulate elastic membrane reshaping in cellular environments

We briefly summarize our approach to develop the computational framework discussed in this work. For simplicity, the governing PDEs considered in our work are restricted to two types – advection-diffusion-reaction (ADR) equations in moving domains (both in the bulk and on the surface) [52, 56] and equations governing Helfrich flow of membranes [39, 57], because they are commonly used to describe the biophysical phenomena underlying changes in cell shape. Our numerical approach consists of the following features:

- Force balance is imposed on surfaces through the use of an adapted version of the algorithm in [48]. The algorithm has been modified to consider the distribution of spontaneous curvature along the geometry, while maintaining the energy-decaying property for Helfrich flows, see Section B.1 for the details. The result is an efficient scheme to simulate membrane deformations starting from realistic resting configurations.
- Mesh update is performed using an Arbitrary Lagrangian-Eulerian (ALE, [58]) two-step approach as introduced in [48] and similar to [55, 59]. As a result, the physical dynamics are decoupled from the geometric update, allowing us to deal with large deformations while maintaining stability. Crucially, the above mentioned energy-decaying property is preserved by the ALE update, as shown in Section B.1. See Section B.2 for additional details.
- ADR systems, either in the bulk or on the surface, are discretized using traditional surface finite element methods [52, 56] and assembled monolithically. As in [55], we adopt the continuous interior penalty (CIP) method to ensure stability of advection-dominated regimes [60] and a structure-preserving scheme to preserve mass and solution bounds [61, 62]. Nonlinearities are dealt with explicitly, see Section B.3 for details.
- Coupling between PDEs is achieved using a staggering approach, as summarized schematically in Figure 1B and detailed in Section 3.

For the full mathematical notation used in this work, see Section A.1 and Figure A.1. In our first example, we simulate migration of cells on flat substrates using coupled surface PDEs on idealized 2D geometries, leading to large deformations. Next, we simulate the extension of chemotactic pseudopods by neutrophils in 3D. Finally, we study the structural plasticity of dendritic spines by considering coupled bulk-surface equations solved for realistic geometries derived from 3D electron microscopy.

### 1.2 Application 1: Mesh redistribution and bounds preservation. Cell motility is regulated by tension-mediated feedback

In our first application, we investigate how ADR equations confined to the boundary of a portion of the cell membrane affect the change in cell shape. This example was chosen to demonstrate the effectiveness of the mesh redistribution algorithm when gradient flow dynamics are absent. We use a model previously published by Lomakin et al. [12] (Figure 2A) to simulate the rearrangement of the actin cytoskeleton along a 1D boundary representing the periphery of the cell-substrate contact region.

**Fig. 2:**
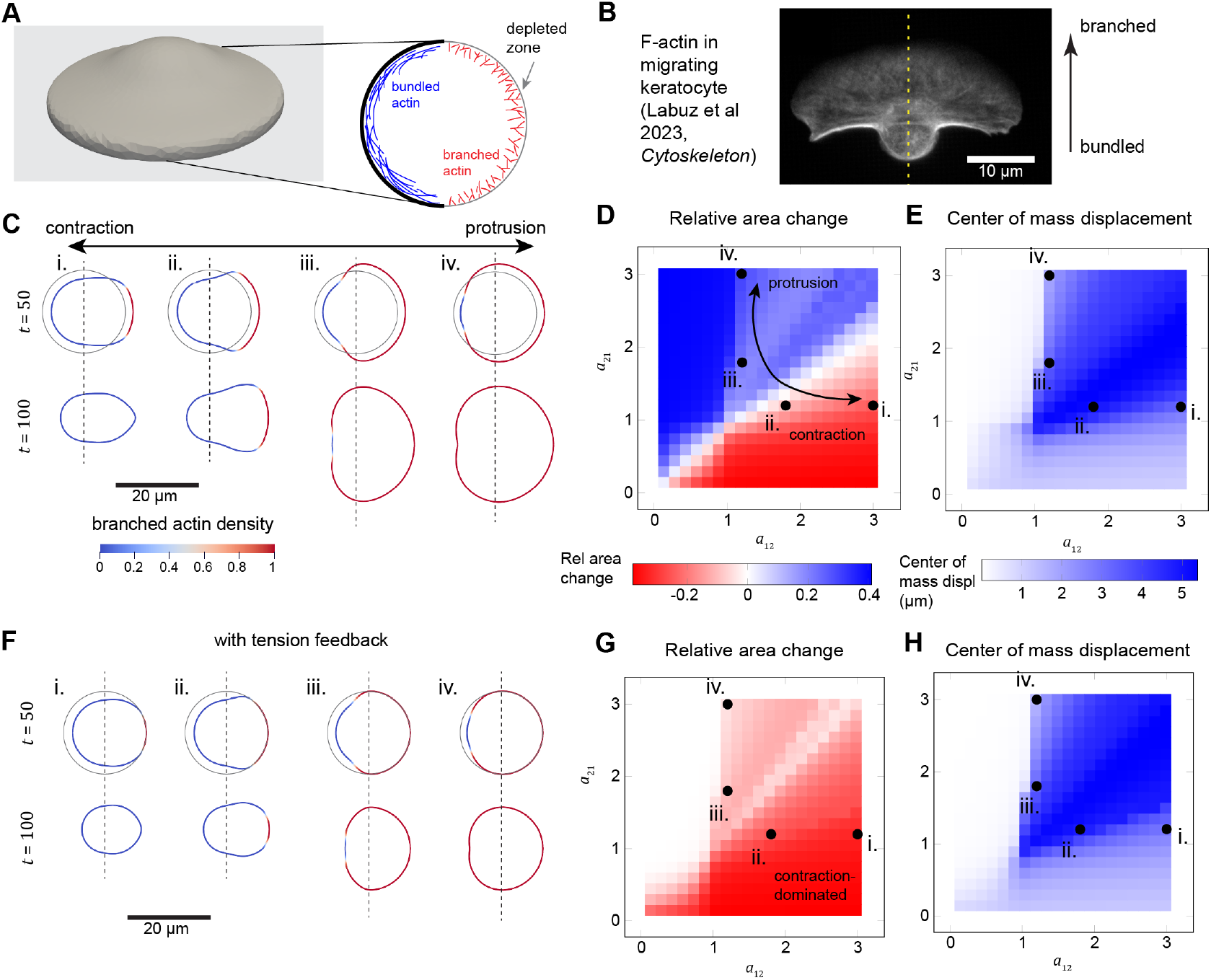
Simulations of cell migration on a 2D substrate reveal the importance of tension feedback. A) Cell geometry on a flat surface, with inset showing the contact region with bundled and branched actin organized along the contact area boundary (contact line). B) Cell morphology and actin localization in a zebrafish keratocyte migrating on a collagen-coated coverslip. Cell was transfected with LifeAct-EGFP for visualization of the cytoskeleton. Image from Fig 1B in [64], reuse authorized by CC-BY license. C) Simulated contact region shapes and density of branched actin for four different combinations of (*a*_12_, *a*_21_) at *t* = 50 s and *t* = 100 s. From left to right, (*a*_12_, *a*_21_) is (1.2, 3), (1.2, 1.8), (1.8, 1.2), and (3, 1.2). Vertical dashed lines indicate the location of the center of mass at *t* = 0. Position of the initial shape is shown as a gray outline for each condition at the 50 s time point. D) Area change |Γ(100)| − |Γ(0)| as a function of (*a*_12_, *a*_21_) with cases shown in panel C indicated by points here. E) Center of mass displacement as a function of (*a*_12_, *a*_21_), computed as 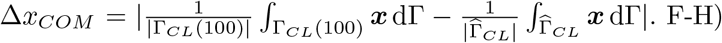 Simulation results with membrane tension feedback, displayed as in C-E. Heatmaps were generated for the combinations (*a*_12_, *a*_21_) = [0.15, 0.30, …, 3] ⊗ [0.15, 0.30, …, 3].

The governing equations of the model [12] consider two biochemical species that represent the density of protrusive branched actin *u*_*A*_ and contractile bundled actin *u*_*B*_ at the boundary of the cell-substrate contact region. The contact region itself is referred to as Ω_*contact*_, whereas its boundary (the contact line) is referred to as Γ_*CL*_. The associated ADR equation for each species on Γ_*CL*_ is given by

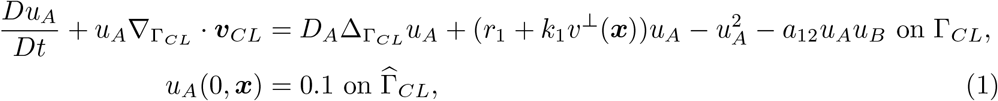

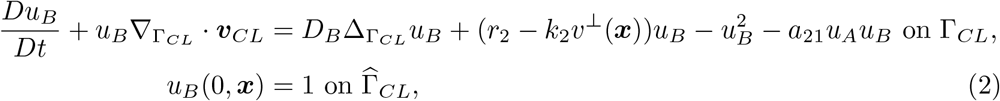

where 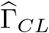 is the contact line domain at 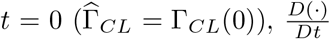 is the material derivative (see Section A), *D*_*A*_ and *D*_*B*_ are diffusion coefficients for *u*_*A*_ and *u*_*B*_ respectively, and *r*_1_, *r*_2_, *k*_1_, and *k*_2_ are kinetic constants defined in Table E3. The coefficient *a*_12_ is the rate of actin bundling and *a*_21_ is the rate of centripetal flow of bundled filaments. The velocity of the contact line, ***v***_*CL*_, is oriented in the normal direction, i.e. 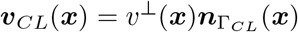, where:

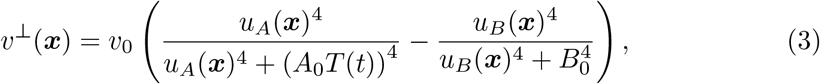

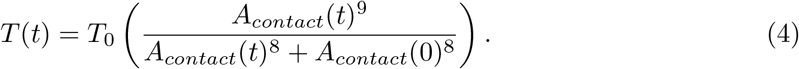

*A*_0_ and *B*_0_ are characteristic concentrations controlling membrane protrusion versus retraction. The time-varying function *T* is the membrane tension, which is an increasing function of membrane contact area 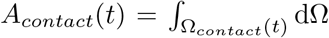. This membrane tension has a stabilizing effect, suppressing radial expansion for large deformations.

The initial contact region is assumed to be circular with radius *R*_cell_. A representation of the computational setup is given in Figure 2A, and the parameter values and descriptions are given in Table E3. Additional details about the algorithm and results from numerical testing can be found in Section C.1 and Figures C.1 and C.7. To ensure stability, the bound- and mass-preserving algorithm proposed in [61, 62] and adopted by the authors in [55] is used to enforce *u*_*A*_, *u*_*B*_ ≥ 0. The mesh redistribution algorithm (Section B.2) is also used to maintain mesh quality and avoid instabilities that arise due to large deformations. In particular, we utilize the Minimal Deformation Rate (MDR) strategy proposed in [63] and adopted in [48, 55]. To understand the role of tension on cell migration, we investigate the case *T* (*t*) ≡ 1 (i.e., no tension feedback) and the case in which tension feedback is present in the form of Equation (4). Just after initializing each simulation, the symmetry is broken by depleting levels of bundled actin over one half of the cell contour as shown in Figure 2A and explained in Section C.1.

Our results without tension feedback are consistent with those established by Lomakin et al. [12]. In particular, the threshold governing transition from cell contraction to expansion of the cell contact region is dictated by the balance between the bundling rate (*a*_12_) and the removal of bundled filaments due to centripetal flow (controlled by *a*_21_). Accordingly, when *a*_12_ *> a*_21_, contraction dominates and the contact region shrinks (Figure 2C-D). Conversely, when *a*_12_ *< a*_21_, protrusion via branched actin dominates and the contact area expands (Figure 2C-D). Motility, as indicated by the movement of the center of mass of the contact region, occurs only when *a*_12_ and *a*_21_ are close in value, resulting in a productive interplay between contraction of the cell rear and expansion of the front (Figure 2C,E). Contact region shapes predicted by our model are similar to those exhibited by migrating cells; for instance, when protrusion slightly dominates over contraction (case iii in Figure 2C), the shape matches that of a migrating keratocyte as shown in Figure 2B from [64]. Furthermore, the model predicts that branched actin filaments dominate towards the wide cell front, whereas bundled contractile filaments accumulate in the cell rear, in good agreement with the fluorescence measurements of actin in Figure 2B.

When we include tension-mediated feedback, we find that the contact area contracts in almost all cases (Figure 2F-H). Tension-mediated feedback eliminates cases of global protrusion observed in the previous case for higher values of *a*_12_, instead favoring a stationary contact area morphology. However, effective motility is still observed when *a*_12_ ≈ *a*_21_ (Figure 2H), where local accumulation of branched actin is sufficient to propel forward motion. Therefore, these simulations predict that membrane tension coordinates cell movement and suppresses global cell deformations, in agreement with the well-accepted idea that membrane tension acts as a global integrator of mechanochemical responses in cell physiology [65–67].

### 1.3 Application 2: Clamped elastic boundaries. Chemotactic protrusions by neutrophils

Next, we model a case of cellular protrusion in 3D driven by biochemical species within the cytosol. As this process is especially amenable to approximation by idealized geometries, we consider chemotactic protrusion by human neutrophils held out of contact with a surface (e.g., in a glass micropipette) and stimulated by a chemoattractant gradient. These neutrophils start from a spherical geometry with radius *R*_*PMN*_ and exhibit isolated protrusions previously referred to as “pure chemotaxis” [68–70]. Different chemoattractant sources have been considered in experiments, including pathogenic particles [70–72] or, as shown here, direct release of chemoattractant molecules from an opposing micropipette [68, 73] (Figure 3A). Here, we consider a minimal computational model for pure chemotaxis, simplified from a previous treatment by Herant et al. [74] (Figure 3A). This problem is chosen to illustrate a new set of numerical challenges beyond the previous example, now involving a 3D interaction between a surface-bulk ADR coupling and a clamped surface moving under Helfrich flow.

**Fig. 3:**
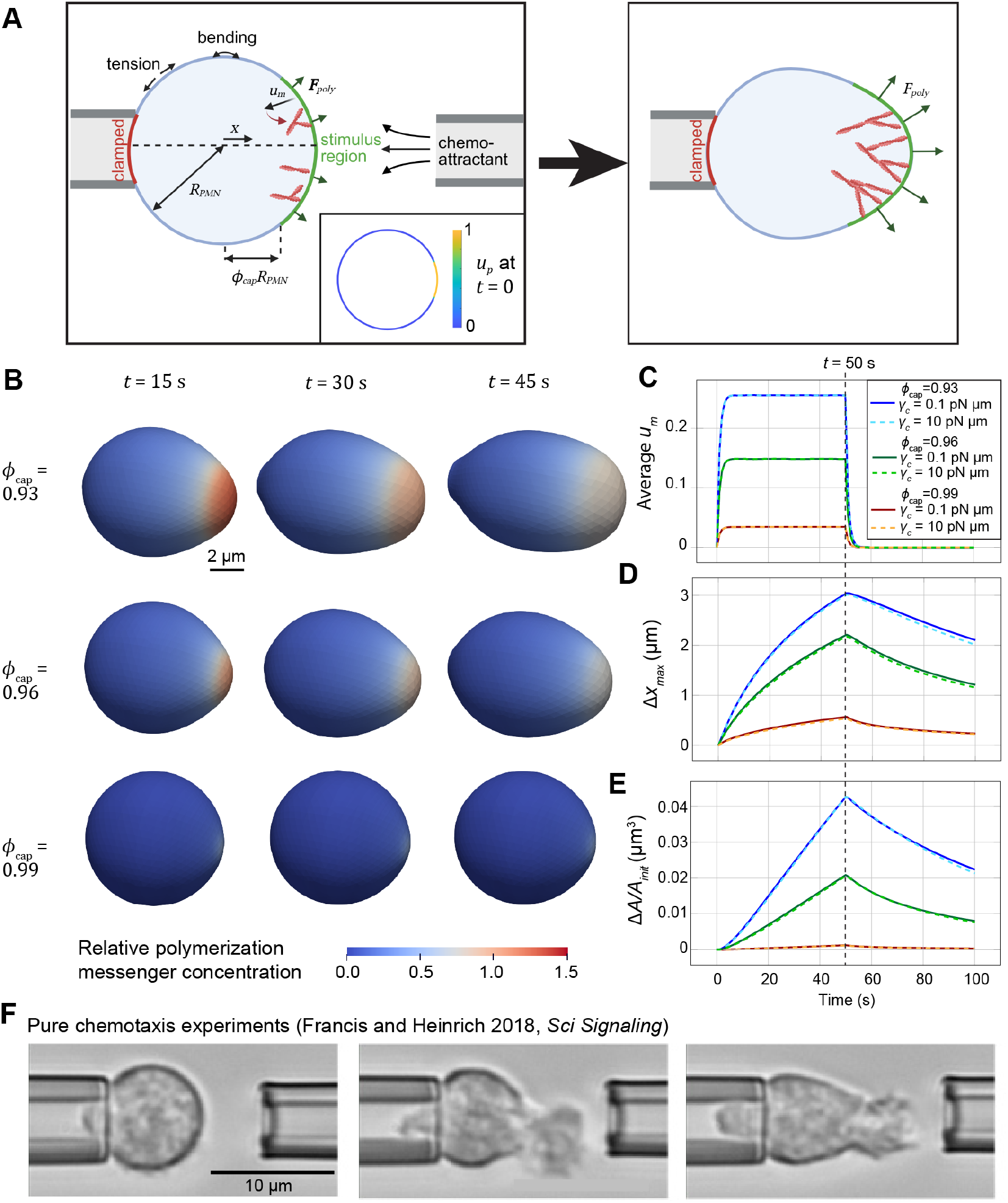
Simulations of pure chemotactic protrusion by human neutrophils. A) Schematic describing the geometry and model components for neutrophil chemotactic protrusion. The initially spherical cell is clamped on the left by a micropipette and deformation is driven by production of *m* over the stimulus region, whose production is restricted by the function *u*_*p*_ in Equation (6), plotted at *t* = 0 in the inset. Sufficiently high *u*_*m*_ is assumed to lead to actin-mediated protrusion, facilitating extension towards the right as shown. Schematic made with Biorender.com. B) Sample images of 3D cell deformations in response to stimulus acting over different spans of the plasma membrane for *γ*_*c*_ = 10.0 pN µm. C-E) Concentration of chemical messenger (C), cell extension in the *x* direction (D), and area expansion (E) for different values of *ϕ*_*cap*_ and *γ*_*c*_. F) Example of human neutrophil morphologies during pure chemotaxis from Fig 3A in Francis and Heinrich 2018 [68], reproduced with permission from The American Association for the Advancement of Science.

In this model, protrusion is driven by the production of a chemical messenger *m* at the plasma membrane Γ_*PM*_, with concentration *u*_*m*_ within the cytosol Ω_*cyto*_, with the governing equation given by

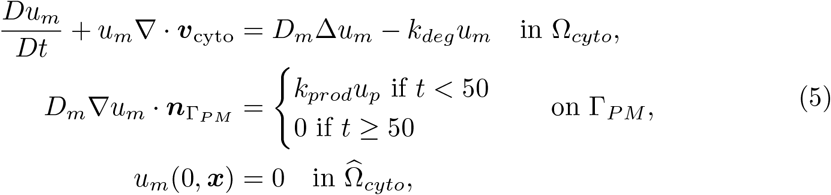

where 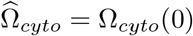, *D*_*m*_ is the diffusion coefficient for *m*, ***v***_*cyto*_ is the extension of the boundary velocity (***v***_*PM*_, defined below) into the cytosol, and *k*_*prod*_ and *k*_*deg*_ are its rates of production at the membrane and degradation in the cytosol, respectively. The production of *u*_*m*_ is restricted to a small initially spherical cap of the membrane over the time interval [0, 50]. The spatial extent of this region was dictated by *ϕ*_*cap*_ using a soft Heaviside function whose transition occurs at *R*_*PMN*_ *ϕ*_*cap*_, as defined in Equation (6) and shown in Figure 3A. We restrict production to the material patch rather than a region with fixed radius. This is achieved by introducing an indicator function *u*_*p*_ that activates production on Γ_*PM*_ . The activation function *u*_*p*_ is modeled through a pure transport equation evolving with the law

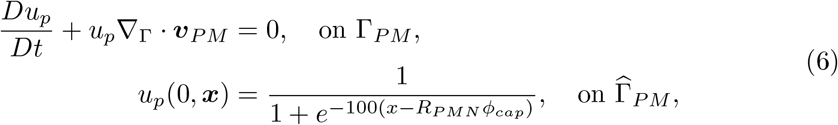

where ***x*** = (*x, y, z*) and 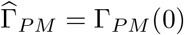. Note that, by following the material evolution of the protruding edge, the activation function maintains the total integrated influx constant during the deformation. Biologically, this variable represents the density of membrane proteins that bind chemoattractant and transduce the signal to the cytosol.

The velocity of the membrane ***v***_*PM*_ results from the local force balance

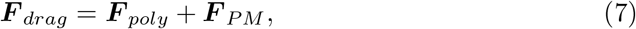

where:

- ***F*** _*drag*_ = *γ*_*drag*_***v***_*PM*_ is the drag force in the overdamped limit between the membrane motion and the extracellular medium;
- 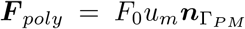 is the local polymerization stress, which is assumed to be directly proportional to *u*_*m*_ and normal to the membrane;
- ***F*** _*PM*_ is the response of the membrane and is modeled using a modified Helfrich functional that accounts for both bending and stretching energies, i.e.:

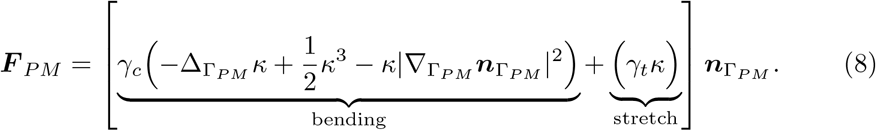

A Lagrange multiplier was also added to preserve volume, see Section 3.1.

The values of the parameters and their descriptions can be found in Table E4, while more details about the algorithm and the results of numerical testing can be found in Section C.2 and Figures C.2, C.4 and C.7.

These simulations predict wide extensions of the neutrophil membrane to the right, towards the presumed location of the chemoattractant source (Figure 3B-D). The extent of deformation depends sensitively on the span of membrane experiencing stimulus. When the stimulus acts over a 43° arc (*ϕ*_cap_ = 0.93), large amounts of *m* are produced, resulting in the formation of a pseudopod matching the overall size of the cell. However, when stimulus is applied over a small region of membrane (about 16°, *ϕ*_cap_ = 0.99), only a small amount of *m* is produced, resulting in a very small local deformation. The computed cell morphologies are similar to those observed experimentally [68]. In particular, larger cell extensions in the lateral direction result in an overall reduction in the cell height in both the model and experiments (Figure 3C,E) as a natural consequence of conservation of cell volume over time. Cell deformations are only slightly affected by the membrane bending modulus *γ*_*c*_, indicating the simulations are conducted in a membrane tension-dominated regime.

In addition to *u*_*m*_ and cell extension (Figure 3C-D), we considered the cell surface area as a key quantity of interest (Figure 3E). Although cell volume is conserved in these simulations, the apparent surface area of cells increases in agreement with previous experiments [20, 68, 71, 75]. The observed deviations in cell surface area (up to about 4%) fall well within the range of deformations supported by unfolding of wrinkles within the neutrophil membrane [75], suggesting that these deformations can occur with minimal increases in membrane tension. Overall, this application showcases the ability of our framework to simulate bulk-surface-reaction-driven deformations of the cell membrane. This model considers a single species, effectively representing the complex network of surface cues integrated at the plasma membrane [4–6], but the computational framework can be extended to multiple species as we demonstrate next.

### 1.4 Application 3: Coupled multiphysics with realistic geometries. Structural plasticity in synapses

In our final example, we demonstrate the power of our framework to solve a nonlinear system of advection-dominant equations in the bulk with an open membrane that undergoes large deformations modeled by Helfrich flow in a realistic dendritic spine geometry (Figure 4). Dendritic spines have particularly complex shapes because of how they form synapses and wrap around axons [76–78]. Many studies have also shown that the shape of dendritic spines is very closely tied to their function and, in particular, to their information-processing capabilities [76, 78, 79]. To test the role of detailed dendritic spine geometries in our simulations, we use a geometry previously considered as an example for adaptive meshing via GAMer2 [38] and reaction-diffusion simulations via SMART [36]. This geometry was derived from volume electron microscopy images in [80] as shown in Figure 4B.

**Fig. 4:**
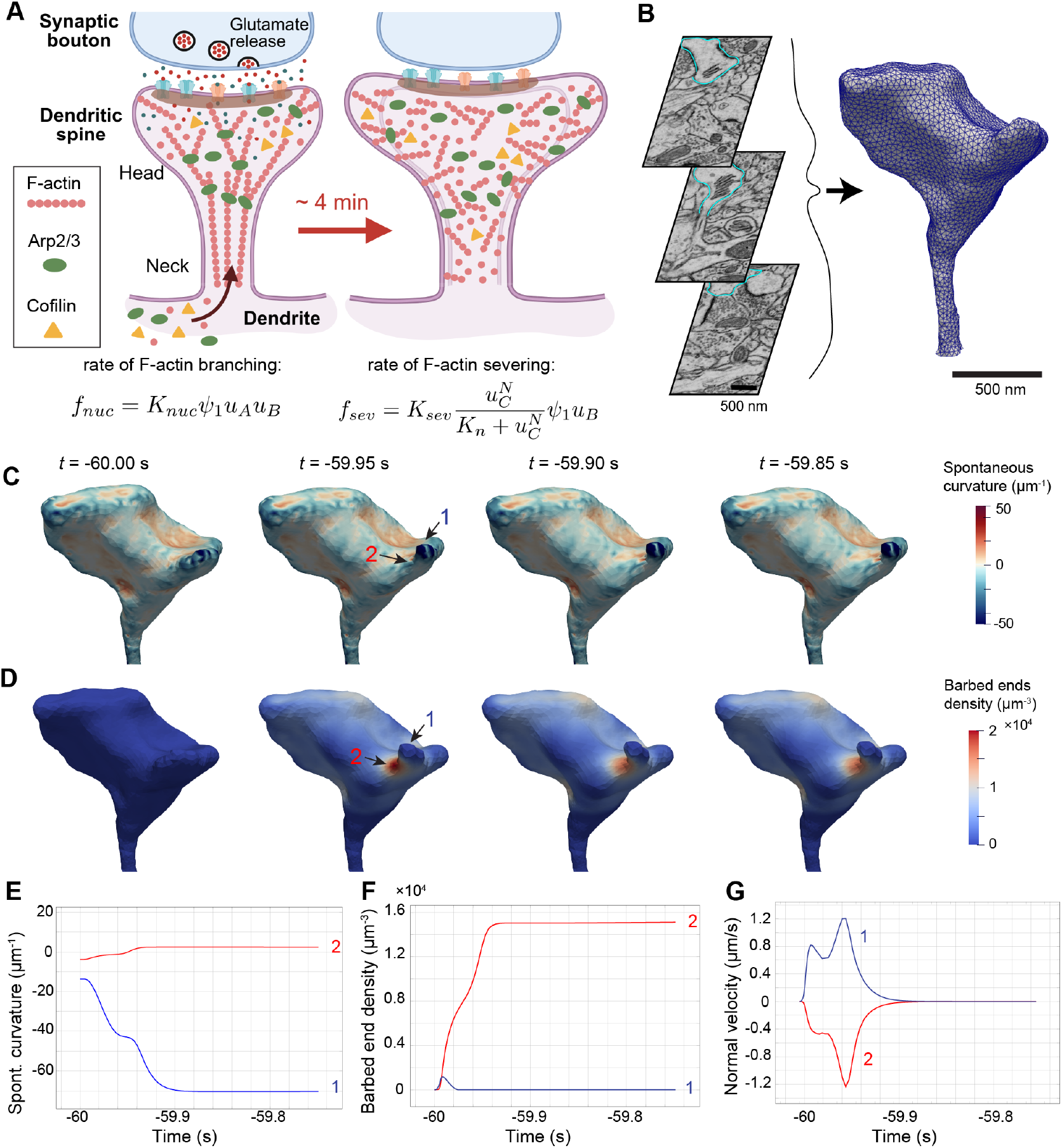
Model of dendritic spine shape changes in a realistic geometry. A) Schematic summarizing model of dendritic spine plasticity, adapted from [81] (reuse authorized by CC-BY license). Glutamate release from the presynaptic cell triggers activation of the dendritic spine, leading to actin assembly and recruitment of ABPs Arp2/3 and cofilin and to subsequent spine swelling. The equations shown govern the reaction rates associated with Arp2/3-mediated branching of actin filaments (left) and cofilin-mediated actin severing (right). Diagram made with Biorender.com. B) Mesh of a dendritic spine segmented from [80] and conditioned in [38], with image slices shown from original dataset (modified from Fig 1A in [38], CC-BY license). C-D) Density of barbed ends (*u*_*B*_; panel C) and spontaneous curvature (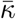 ;panel D) plotted over the realistic spine mesh for the first 0.15 s of simulation. Point 1 denotes the region of most rapid outward membrane movement and point 2 is a location at the base of this expanding region. E-G) Plots showing the spontaneous curvature (E), barbed end density (F), and normal velocity of the membrane (G) at points of interest during the initial 0.25 s of simulation.

Building on the model of Bonilla-Quintana et al. [81], we simulate the dramatic volumetric growth of a dendritic spine in response to actin-mediated forces when stimulated at the synaptic terminal (Figure 4A). In this model, deformation is driven by the concentration of actin barbed ends per unit volume, *u*_*B*_. The localization of barbed ends is modulated by actin-binding proteins (ABPs) including Arp2/3 (concentration *u*_*A*_), which facilitates actin branching, and cofilin (concentration *u*_*C*_), which severs actin filaments. The governing equations for these three species are given by

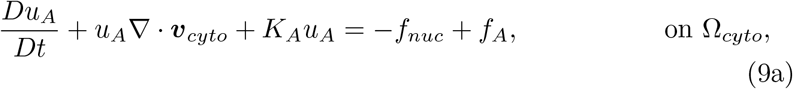

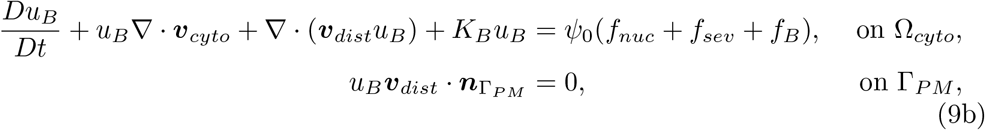

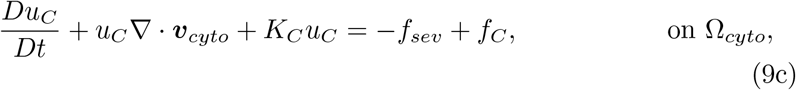

where *K*_*A*_, *K*_*B*_, and *K*_*C*_ are degradation rates for each species and *ψ*_0_ is a unit conversion factor (see Table E5). The velocity ***v***_*cyto*_ represents the advective flow of species within the cytosol and ***v***_*dist*_ serves as a proxy to simulate the growth of the branched actin towards the membrane. Nonlinear chemical interactions associated with the ABPs (*f*_*nuc*_ and *f*_*sev*_) are modeled as in [81], as shown in Figure 4A. In addition, a constant basal influx and an impulse are modeled as *f*_*i*_ = *I*_*i*_ + *I*_*S,i*_(*t*), *i* ∈ {*A, B, C*} ; in contrast to [81], these source terms act on the entire domain rather than being confined to the spine head. The impulse function is a phenomenological representation of the complex biochemical events underlying structural plasticity [76, 82, 83].

Similar to [81], the velocity ***v***_*dist*_ is related to the signed distance function *d*_*s*_ associated with Γ_*PM*_, which in this article is approximated using the heat method [84]. Rather than simply defining the velocity to be proportional to the gradient of *d*_*s*_ as in [81], we assign:

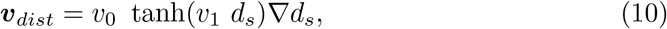

which ensures that the relative velocity goes to zero at the membrane for physical consistency. In order to avoid instabilities, the positivity and bound-preserving algorithm proposed in [61, 62] and adopted by the authors in [55] has been used to keep the concentrations of *u*_*A*_, *u*_*B*_, *u*_*C*_ ≥ 0. As in [81], the impulses *I*_*S,i*_(*t*) take the form

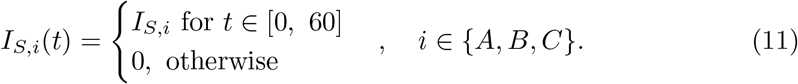

The principle underlying the force balance at the membrane is analogous to the previous example in Equation (7):

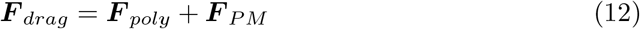

where, once again, ***F*** _*drag*_ = *γ*_*drag*_***v***_*PM*_ . The polymerization stress is now determined by the density of barbed ends; that is, 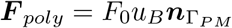. Defining ***F*** _*PM*_ for this case requires additional care, because the dendritic spine geometry is considerably more complex and Helfrich flow with constant spontaneous curvature 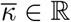 does not yield a stationary solution in the absence of external forces. Accordingly, we consider generalized energies where 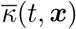 is heterogeneous and assume that 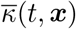 is transported with the deforming membrane. Mathematically, this is encoded by assuming that the normal time derivative of the spontaneous curvature is zero (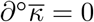, Section A, C.3). The resulting scheme is energy-decaying regardless of the initial approximation of the mean curvature and Weingarten map (i.e., ∇_Γ_***n***_Γ_). The membrane stress now assumes the form

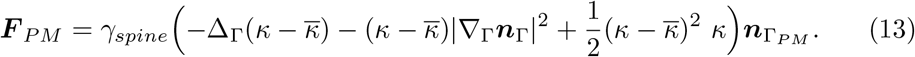

The resulting velocity ***v***_*PM*_ is extended into the cytosol to yield ***v***_*cyto*_, as described in Section B.2. All parameter values and descriptions can be found in Table E5. More details about the overall algorithm can be found in Section B.1, and results of tests that prove the stability of this scheme can be found in Section C.3 with associated figures Figures C.5 to C.7.

We now test the predictions of our full model with coupled ADR and Helfrich flow. Upon initializing the simulation, but prior to adding additional stimulus (*t <* 0), we observe an initial rapid change in the spine morphology. Specifically, one highly negatively curved region of the shape immediately protrudes outward upon introducing barbed ends into the system (point 1 in Figure 4). This protrusion is driven by the negative spontaneous curvature in that region, as shown in Figure 4C,E. This leads to the depletion of barbed ends within this expanding region, but an accumulation at the base of the extension (point 2 in Figure 4) where inward movement of the membrane dominates (Figure 4D-E). The inverse correlation between actin density and outward expansion differs from previous examples. This is a consequence of the outward membrane movement locally increasing cytosolic volume and thereby diluting the barbed ends. Conversely, the constant flow of barbed ends to the cell periphery results in regions of aggregation where the membrane locally retracts. This rapid early expansion was not observed in simulations with an idealized spine geometry (Figure D.1), underscoring the importance of considering realistic geometries.

We next simulate the growth of the spine in response to the events that induce structural plasticity [81, 85]. Prior to additional stimulus that is associated with longterm potentiation at *t* = 0, we observe expansion of the spine head during the approach to a new steady-state. During this phase, regions dense with barbed ends (points D1 and D2 in Figure 5) are also associated with inward membrane movement (Figure 5E-F), in agreement with Figure 4. Regions with sparse concentrations of barbed ends (points S1 and S2 in Figure 5) are associated with outward membrane movement. Upon providing additional stimulus, the average concentration of barbed ends throughout the spine increases faster, resulting in fast growth of the spine head radius as anticipated (Figure 5C-G). The spine neck radius remains largely unchanged throughout the simulation (Figure 5G). This result is consistent with previous model predictions on spine volume changes observed in structural plasticity [81]. We note that the growth of the spine head radius does not scale directly with the increase in barbed ends throughout the spine head. Rather, growth appears to stall despite a rapid accumulation of barbed ends throughout the spine head. Here, the negative feedback arises from membrane bending, which is a different mechanism from membrane tension-mediated mechanical feedback in the previous examples and in the literature [66, 67]. We also find that the overall membrane displacement depends on the initial curvature in a given region. In particular, we find that regions that were initially flatter exhibited larger amounts of expansion (Figure 5C, Figure D.2). Overall, our approach shows the ability to make several nontrivial predictions that depend on localized mechanochemical feedback.

**Fig. 5:**
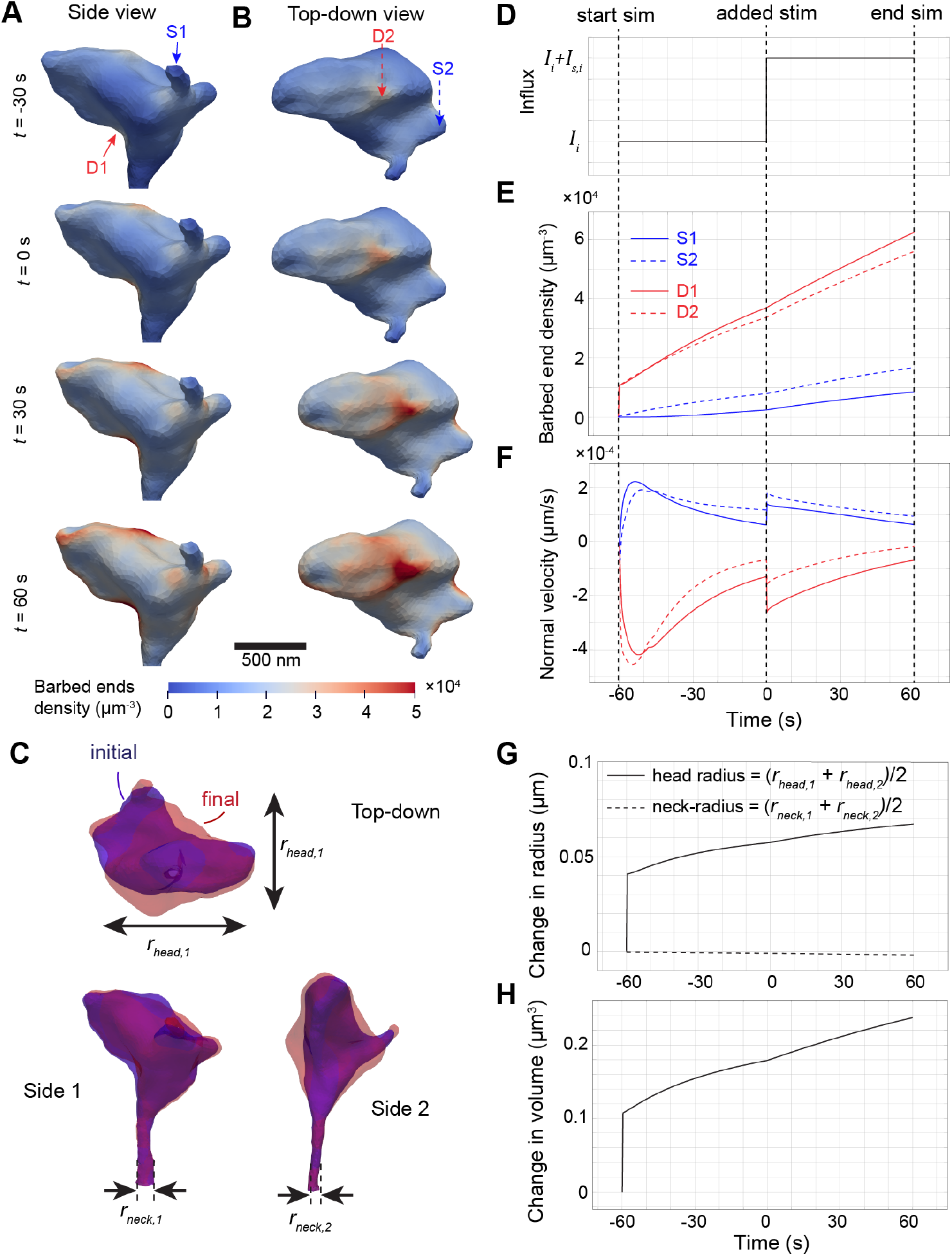
Simulation of dendritic spine shape changes. A-B) Side view (A) and top-down view (B) of spine expansion and barbed end density over the simulation. S1 and S2 denote regions of sparse barbed ends, whereas D1 and D2 denote regions of dense barbed ends. C) Initial (blue) and final (red) configurations of the dendritic spine from different viewpoints, including definitions of the effective radii of the spine head and spine neck. D-F) Plots showing the time-dependent stimulus function (D) and the resulting changes to barbed end density (E) and normal membrane velocity (F) at each point of interest. G-H) Effective change in spine head radius and spine neck radius over time (G), leading to changes in global spine volume (H).

## 2 Discussion

In this study, we presented a finite-element-based simulation framework to simulate moving boundary problems in cell biophysics, including equations governing membrane kinematics and those governing the reaction and transport of biochemical species. We chose three applications with sufficient complexity to illustrate the range of problems that we can readily simulate. The first application considered an example in which membrane velocity is directly dictated by the concentration of species on the boundary of a two-dimensional shape. The two additional applications treat the cell membrane as an elastic boundary, assuming that it follows a gradient flow to minimize the Helfrich potential energy. In all cases, mechanics and geometry are tightly coupled to a biochemical ADR system, presenting a nontrivial numerical challenge. We demonstrate that our methods provide mathematically robust and flexible strategies to simulate these models that were solved using different numerical approaches across the literature including FEM [74], the finite difference method [81], and graded radial extension [12]. Our approach complements the strengths of previous treatments by allowing us to enforce important biophysical constraints such as conservation of mass and non-negativity of biochemical species concentrations and to consider detailed realistic cellular geometries.

It is increasingly appreciated that models of biochemical reaction and transport often yield characteristically different predictions in realistic cellular geometries compared to their idealized counterparts [36, 77, 86]. However, most simulations in realistic geometries are restricted to stationary geometries. By allowing for a nonuniform spontaneous curvature that is transported with the geometry during cell deformation, our approach lets us systematically explore deviations from a stable biological morphology measured by high-resolution microscopy and make biophysically founded predictions that may not be intuitively obvious. For example, the rapid initial deformation exhibited by a particular region of the realistic dendritic spine in our final application was a new phenomenon not previously identified when the same model was considered within idealized spines [81]. Accordingly, the highly nontrivial shapes of biomembranes, both at the cell boundary and within intracellular organelles, necessitate new tools such as ours capable of generating biophysical predictions with a new level of realism.

The unexpected nature of some model predictions arises not just from the complicated geometry, but from the nonlinear bidirectional coupling between membrane mechanics and biochemical reaction-transport. In particular, the rapid expansion and contraction of different regions of the dendritic spine head reflect the sensitivity of mechanochemical coupling to the local curvature of the cell membrane. Coupling between local curvature and bulk-surface reactions also plays an important role in the model of chemotactic protrusion considered here. Generally, coupling between cell shape and biochemical reactions plays a dominant role in many different biophysical models, such as those for cell spreading [87, 88] and for cell motility [40, 41]. In fact, mechanochemical coupling and membrane-mediated feedback on chemical reactions are likely a universal feature across different cellular and multicellular phenomena. Therefore, our approaches here provide opportunities to explore other applications across scales, extending to multicellular problems in developmental biology and cancer biology.

From a technical perspective, this work builds on a number of approaches, most prominently those developed for the SFEM. It belongs to the family of FEM frameworks built to tackle complex cellular reshaping where membrane kinematics plays a key role [49, 50, 53, 54, 89, 90]. In particular, it builds on recent advances in structure-preserving Helfrich flow discretizations [48, 59, 63] to allow for fully 3D simulations on realistic geometries that are possibly open. It then adopts a post-processing approach to decouple physics evolution from geometric evolution in a consistent way as done in [48]. Finally, it borrows from established coupling approaches [90] and bound-preserving algorithms [61, 62] to approach the complexity of multi-physics phenomena in both the bulk and on the surface in a stable, accurate way.

Numerically, realistic geometries present an additional complexity for initializing the simulations with a membrane bending energy. In fact, what is desired is a true energetic ground state that keeps the shape stable when no external stimulus is present. While this is automatically achieved using idealized geometries with zero or constant spontaneous curvature, such as spheres or tori (Clifford torus), it is not the case in general. We solved this problem by extending the algorithm in [48] to consider a heterogeneous spontaneous curvature (Section B.1). This is closely linked to the assumption that every point in the membrane has a characteristic spontaneous curvature that allows for an initial ground state to exist in the first place. The novel, extended algorithm was able to maintain all the properties of the previous one and unlock stable Helfrich flow simulations on generalized meshes. To the authors’ knowledge, this is the first such result for Helfrich flows within the SFEM framework.

In addition to these new features of our framework, there are many opportunities to integrate this work with techniques developed across computational biophysics. For instance, as the systems of equations grow larger for complicated networks of cell signaling, it will be important to integrate our approaches with fast multipole solvers for PDEs [91]. Furthermore, our current framework does not support topological changes observed during processes such as endocytosis, exocytosis, and phagocytosis; to fully represent such topological changes in membranes, we will need to use other approaches such as phase-field [92, 93], level-set methods [94, 95] or unfitted finite element methods [96, 97]. Finally, given the high levels of uncertainty associated with parameters in biological models, it is crucial to incorporate approaches from uncertainty quantification. This could facilitate parameter estimation algorithms [98, 99] as well as other inverse problems in cell biophysics such as force inference [100, 101].

In summary, this work opens new opportunities for biophysical simulations with higher levels of biochemical and geometric detail. As an open-source toolkit, our code can be readily adapted to various problems and expanded to integrate with other frameworks, allowing us to generate new predictions in the field of computational cell biophysics.

## 3 Methods

We consider a moving surface Γ(*t*) that evolves following the material map **Φ**(*t*, ·) : 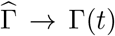, where 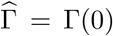 is the initial, reference surface. The map has the associated velocity 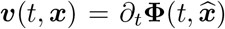 where 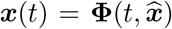. We also consider the ALE map 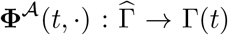 to describe the motion of the mesh points, with associated velocity 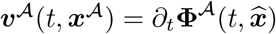, with 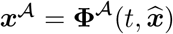. The normal velocity is uniquely determined and the notation *v*^⊥^ is used for it. If a bulk domain is considered, these definitions can be easily adapted (see Section A). The dynamics of the overall system is computed at a number of discrete time points *t*_0_, · · ·, *t*_*N*_ = *t*_*end*_. At each time interval [*t*_*n*_, *t*_*n*+1_] with timestep *τ* = *t*_*n*+1_ − *t*_*n*_, *n* = 0, …, *N* − 1 a sequence of three steps is inserted in a staggering loop. The steps are as follows:

- **Step 1**: A force balance equation is formulated on the surface. Given the problems considered in this work, the balance reduces to an equation for the normal velocity of the surface that assumes the expression *v*^⊥^ = (…). If a tangential component is present, the algorithm can be readily adapted;
- **Step 2**: A geometrical update is performed by moving the surface mesh vertices according to the ALE velocity ***v***^*A*^ and current time step length *τ* . If no ALE map is necessary, the material velocity ***v*** is used instead. If volume elements are present, the ALE motion is extended to the bulk by solving an auxiliary harmonic problem;
- **Step 3**: The ADR system is solved on the updated geometry. The coupling between the ADR system and the geometrical update is nonlinear, as the geometry takes part in the definition of the PDEs and the PDEs take part in the definition of the velocity *v*^⊥^.

Steps 1, 2, and 3 are then inserted in a staggering loop using a fixed-point iteration to solve for the nonlinearities characteristic of the system. In the following sections, we detail the equations considered in the general framework and how we solve them. The basic notation for the continuum quantities is introduced in Appendix A.

### 3.1 Step 1: Surface balance equation

We consider surface motions that are either prescribed or resulting from an elastic response of the membrane to external forces. In both cases, we are interested in computing the normal component of the velocity that governs the geometric evolution of the surface.

Our starting point for the elastic case is the generalized Helfrich-type potential in the form

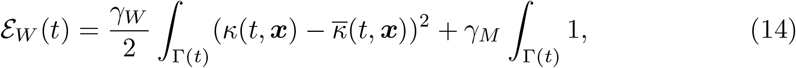

where *κ* is the mean curvature of the surface, 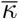 is the spontaneous curvature, which can be heterogeneous in space and time, *γ*_*W*_ is the bending modulus and *γ*_*M*_ is the tension. Computing the variation dℰ_*W*_ */* d*t* [102] under the hypothesis 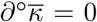 leads to the following equation for the normal velocity *v*^⊥^ = ***v***^⊥^ · ***n***_Γ_ of the surface:

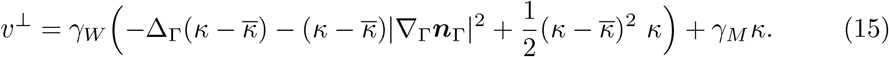

External influences can thus be incorporated as an additional right-hand side *v*^⊥^ = … + *f*, similar to what has already been done in the literature, see [51] and references therein. Moreover, additional flows (mean curvature flow, surface diffusion flow) and constraints (area preservation, volume preservation) can be directly added to this formula. Following the advances presented in [48] and our proposed extension to account for heterogeneous spontaneous curvature (see Section B.1), this fourth order equation can be closed by adding additional equations for the mean curvature *κ* and the spontaneous curvature 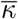. These read:

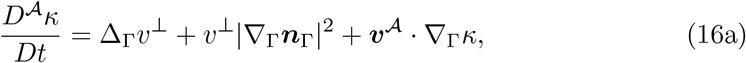

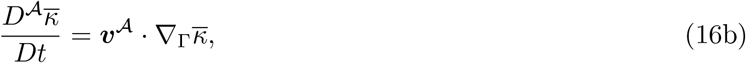

where 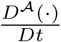 is the total ALE derivative and ***v***^*A*^ is the ALE velocity of the surface (see Section A). Equation (15) together with Equation (16b) constitute our basis equation system governing membrane motion.

For the weak formulation, space-time discretization and properties of this balance equation, including boundary conditions, we refer to Section B.1.

### 3.2 Step 2: Geometric update

Once the surface velocity is known, the geometry is evolved using the two-stage procedure introduced in [48]. The surface ALE parametrization 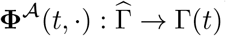 evolving the reference surface 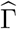 is required to satisfy the system

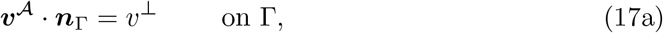

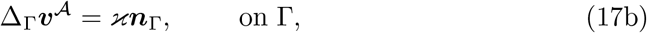

where ϰ is a Lagrange multiplier. In practice, the normal ALE velocity has to agree with the normal velocity prescribed by the force balance in Equation (17a). The tangential ALE motion, on the other hand, obeys Equation (17b), which implements one of the many redistribution strategies proposed to preserve mesh quality along the simulation [63]. An alternative choice would be to substitute Equation (17b) with Δ_Γ_**Φ**^*A*^ = ϰ***n***_Γ_, which is the preferred choice in [48].

Once the map has been computed on the surface, an extension might be needed for the bulk. In this case, we choose to prolong the map into the volume by solving a simple harmonic problem:

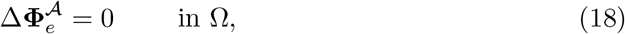

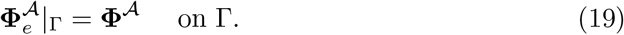

Notice that with the ALE maps used here, the definition of **Φ**^*A*^, 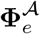 only depends on the normal velocity *v*^⊥^ and not on the material tangential velocity ***v***^∥^. This is because the geometric shape of the domain is uniquely determined by *v*^⊥^ and it is also the reason why we only considered *v*^⊥^ in Section 3.1. This is in line with the idea of an ALE approach where the normal motion is encoded in Lagrangian terms, while the tangential motion can be encoded in mixed Lagrangian–Eulerian terms.

For the weak formulation, space-time discretization and properties of this geometric update we refer to Section B.2.

*Remark 1* When the material velocity ***v*** is unknown in the bulk, we make the additional choice to extend the surface material velocity ***v*** = *v*^⊥^***n***_Γ_ using the same harmonic approach presented above. Although this might not be physically motivated, it gives us a simple way of creating a mass-preserving evolving-domain simulation for the species considered. In Section 1.3, we expect such an advection field to be negligible with respect to the diffusion, while in Section 1.4 such a choice allows for exact zero-flux boundary conditions.

### 3.3 Step 3: Advection-diffusion-reaction equations

The model system of surface-bulk ADR equations we consider reads:

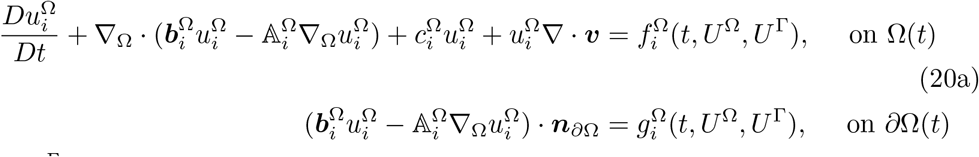

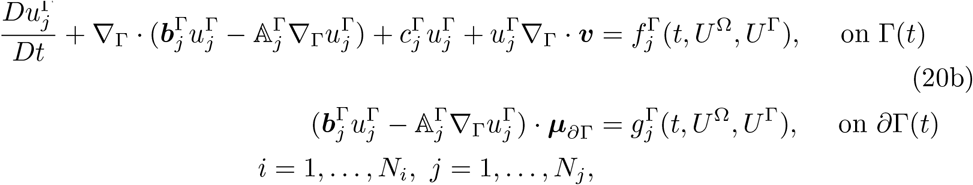

where ***v*** is the unique material velocity, 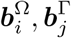 are the relative advection fields, 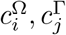 are reaction rates, 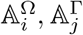 are diffusion matrices. The right-hand sides 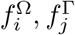 group additional contributions that can potentially depend nonlinearly on the collection of species in the bulk 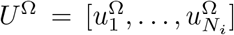 and on the surface 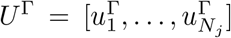. The same holds for 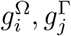, which describe boundary conditions. For compatibility conditions, 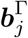 have to be tangential vector fields and 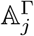 have to map tangential vector fields into tangential vector fields.

The form chosen allows for a common weak formulation for any species *u*_*i*_ in the system that reads: find 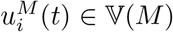 such that

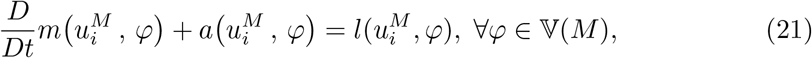

where

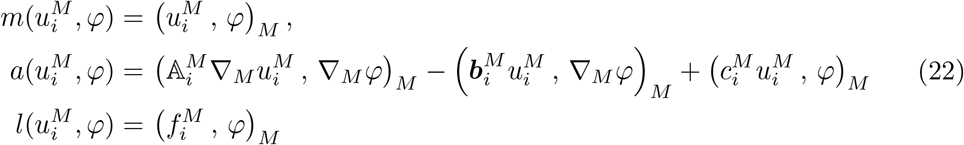

and *M* ∈ {Ω(*t*), Γ(*t*)} is the domain of definition. Equation (22) results from imposing homogeneous Neumann boundary conditions on the entire *∂M*, an assumption for which the appropriate function space is V(*M*) := *H*^1^(*M*) = *W* ^1,2^(*M*), i.e. the order one Sobolev space of square-integrable functions (see Section A). The bilinear forms and function space can be adapted to consider different boundary conditions, see Section B.3. In the above, we used the material map **Φ** and the material velocity ***v***. If an ALE approach is used, it can be shown (see Section A) that it is sufficient to add an additional relative velocity ***w***(*t*, ***x***) = ***v***(*t*, ***x***) − ***v***^*A*^(*t*, ***x***) ∈ *T*_***x***_*M* (*t*).

### 3.4 Coupling strategy

The staggering loop for Steps 1 through 3 is based on a fixed-point approach (see Figure 1). Initial values required for a general internal loop at time *t*_*n*_ are:

- Previous and current geometrical configurations 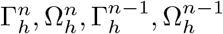.
- The current timestep *τ*^*n*^, which also defines the timestep interval [*t*_*n*_, *t*_*n*+1_] where *t*_*n*+1_ = *t*_*n*_ + *τ*^*n*^.
- The solutions of the solvers at *t*_*n*_, which here will be generally indicated as the solution vector 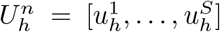 with *S* the number of solvers. Note that 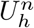 includes the solution of the ALE solver, and thus discrete functions encoding the domain displacement and velocity.
- Minimum and maximum tolerances for the loop {*tol*_*min*_, *tol*_*max*_} together with a maximum number of allowed iterations count *iter*_*max*_.
- Two solution vectors 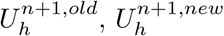 used to check convergence tolerances for the new timestep. We set 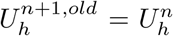 and 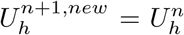 at the beginning of the loop.

The following loop is then executed for a maximum of *iter*_*max*_ times:

- Solve Step 1 using the values in 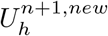 if needed, then update 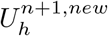 with the new solutions.
- Solve Step 2 using the values in 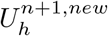 if needed, then update 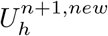 with the new solutions.
- Solve Step 3 using the values in 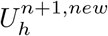 if needed, then update 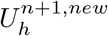 with the new solutions.
- Compute per-solver tolerances *tol*_*i*_, *i* = 1, …, *S* for the difference between 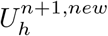 and 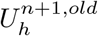 and retain *tol* := max_*i*_ *tol*_*i*_.
- If *tol < tol*_*max*_ the desired accuracy has been reached and the loop is broken. If not, set 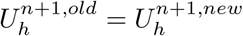 and repeat the loop.

Once the above loop is terminated, the cases are as follows:

- If *tol* ⩽ *tol*_*max*_ then the step is a valid step and we can move forward to *t*_*n*+1_ by setting 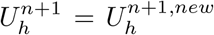 and updating the domain configuration. If an adaptive timestep is used and *tol < tol*_*min*_, then *τ*^*n*+1^ is updated to a higher value *τ*^*n*+1^ = *C*_0_ ∗ *τ*^*n*^ where *C*_0_ *>* 1 is a user-designed scaling factor.
- If *tol > tol*_*max*_ then the step is invalid. If no adaptive strategy is adopted, then an error is thrown. If an adaptive strategy is adopted instead, the solutions of the solvers are reset as 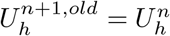 and 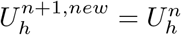, and the timestep is reduced to 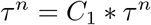 where *C*_1_ *<* 1 is a user-designed scaling factor. After this, the loop is repeated until a valid step is encountered or a minimum value of *τ*^*n*^ is reached, the latter triggering an error.

## Supplementary information

Appendices A-C are included with 8 associated supplementary figures. Appendix D includes 2 additional supplementary figures referenced in the text and Appendix E includes 3 tables stating the parameters used in each of the simulations within this paper.

## Acknowledgements

This work was supported by the National Science Foundation under grant EEC-2127509 to the American Society for Engineering Education (to E.A.F.) and under AFOSR FA9550-25-1-0304 (to P.R.).

## Author contributions statement

A.C. wrote the code, conducted all the simulations and numerical testing, and authored all of the appendices. A.C., E.A.F., A.M., and P.R. drafted and edited the manuscript. E.A.F. and A.C. prepared the figures and analyzed the results. A.M. and P.R. designed and supervised the project and obtained funding. All authors reviewed and approved the final version.

## Declaration of generative AI and AI-assisted technologies in the manuscript preparation process

During the preparation of this work the author(s) used Claude to check spelling and grammar. After using this tool/service, the author(s) reviewed and edited the content as needed and take(s) full responsibility for the content of the published article.

## Appendix A Domain description and space-time discretization

Let *M* ⊂ ℝ^*d*^, *d* = 2, 3, be an *m*-dimensional oriented, compact, *C*^2^ manifold with boundary *∂M* . The notation Ω ≡ *M* when dim (*M*) = *d* and Γ ≡ *M* when dim (*M*) = *d* − 1 will also be used. In the case dim(*M*) = *d* − 1, we denote with *T*_***p***_(*M*) and ***n***_*M*_ (***p***) the tangent and normal space with respect to ***p***, respectively. The *tangential projection* at ***p*** is then defined as

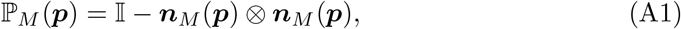

where I is the identity matrix. Differential operators can be defined as follows:

- gradient: 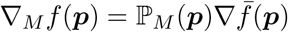
- divergence: ∇_*M*_ · *f* (***p***) = Tr(∇_*M*_ *f* (***p***))
- Laplace-Beltrami operator: Δ_*M*_ *f* (***p***) = ∇_*M*_ · ∇_*M*_ *f* (***p***)

where 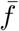 is a smooth extension of *f* to a *d*-dimensional neighborhood of *M* such that 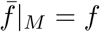 and ∇ is the Euclidean gradient in ℝ^*d*^. For ***f*** : *M* → ℝ^*d*^ and F : *M* → ℝ^*d×d*^ we define (∇_*M*_ ***f***)_*ij*_ = (∇(***f*** · ***e***_*i*_))_*j*_ and 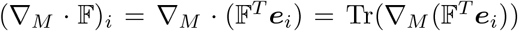, where ***e***_*i*_, *i* ∈ {1, …, *d*} is the canonical basis in ℝ^*d*^. This allows us to define Δ_*M*_ ***f*** = ∇_*M*_ · ∇_*M*_ ***f*** . The curvature properties of *M* are captured by

- the *extended Weingarten map*: 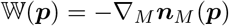;
- the *mean curvature*: *κ*(***p***) = Tr(W(***p***)).

A sketch of possible model setups is shown in Figure A.1.

**Fig. A.1:**
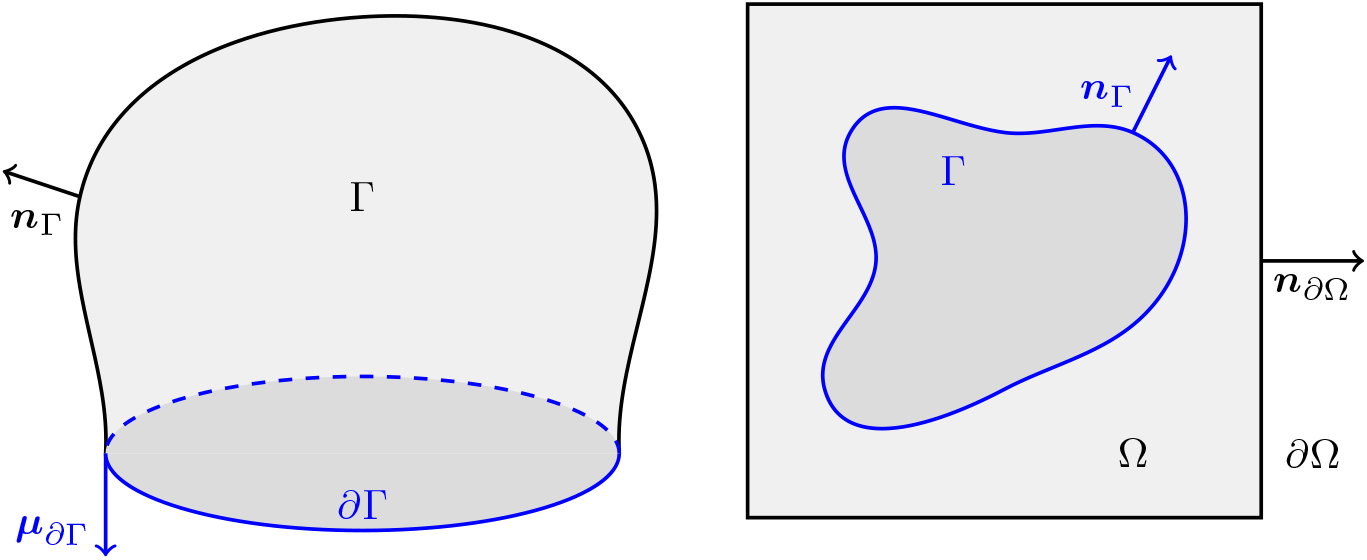
Sketch of an open and closed geometry for notation purposes. One is a 3D object and the other is a 2D object.

We consider a time interval *J* = [0, *t*_*end*_], *t*_*end*_ *>* 0 and a *C*^2^-*evolving manifold* {*M* (*t*)}_*t*∈*J*_ in ℝ^*d*^. In this article an evolving manifold is modeled by a *reference manifold* 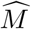 together with a *flow map*

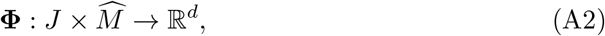

such that

- denoting 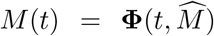, the map 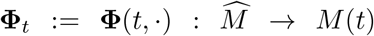 is a *C*^2^ diffeomorphism with inverse map 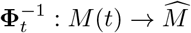;
- 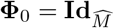 where 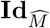 is the identity map on the reference manifold;
- **Φ, Φ** ∈ *C* (*J*; *C* (ℝ^*d*^; ℝ^*d*^)).

The mapping **Φ**_*t*_ can be used to define a pull-forward map of functions [103, 104]

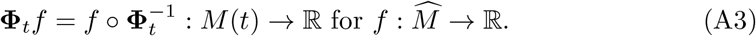

We assume that there exists a velocity field ***v*** : *J ×* ℝ^*d*^ → ℝ^*d*^ with ***v*** ∈ *C*^0^(*J*; *C*^2^(ℝ^*d*^; ℝ^*d*^)) such that for any *t* ∈ *J* and every 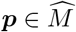

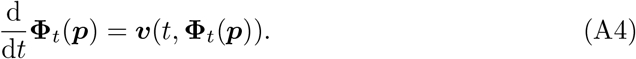

For ***x*** ∈ *M* (*t*), the velocity ***v*** can be split into a tangential component ***v***^∥^(*t*, ***x***) = ℙ_*M*_ (*t*, ***x***)***v***(*t*, ***x***) ∈ *T*_***x***_*M* (*t*) and a normal one ***v***^⊥^(*t*, ***x***) = ***v***(*t*, ***x***) − ***v***^∥^(*t*, ***x***). This allows us to define

- the *normal derivative*: 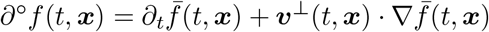;
- the *material derivative*: 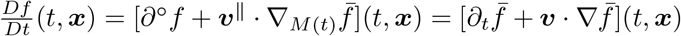.

The definition of a flow map is not unique and the same evolving manifold can be described by different maps. The flow map of the computational domain can be chosen so as to maintain good mesh properties. This is achieved by introducing a second flow map **Φ**^*A*^ called *Arbitrary Lagrangian Eulerian (ALE) map* with corresponding velocity ***v***^*A*^ and ALE material derivative 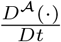. For surfaces, it holds

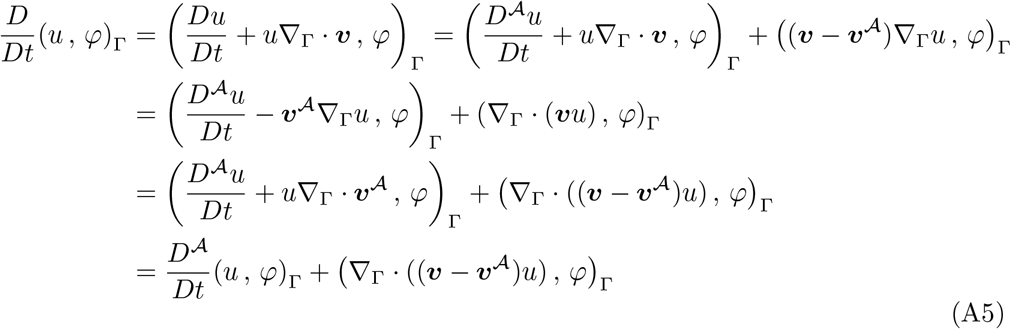

where 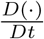 indicates the total derivative on Γ considered as the evolution of the ALE map **Φ**^*A*^ instead of the material map **Φ**. If an ALE approach is used, then one can just add a relative velocity field ***w*** = ***v*** − ***v***^*A*^, which is indeed tangential, and compensate for the difference between the two maps. The relation for the bulk case is analogous and follows simplified arguments. It is worth noting that the normal velocity ***v***^⊥^ is in any case uniquely determined, i.e. ***v***^⊥^(*t*, ·) = ***v***^*A*,⊥^(*t*, ·), ∀*t* ∈ *J*, including the normal velocity of the boundary *∂M* (*t*). From this, it follows that the normal derivative *∂*^°^ is also uniquely determined independently on the parametrization used for the domain evolution.

We use the notation V(*M*) := *H*^1^(*M*) = *W* ^1,2^(*M*) for the order one Sobolev space of square-integrable functions. We also introduce the notation

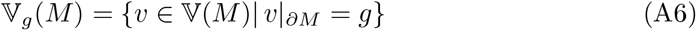

for functions in V(*M*) that assume a user-given value along the boundary of the domain. For a precise definition of such spaces for evolving domains see [103, 104].

### Fig. A.1 Space and time discretization

The domain geometries are discretized using piecewise linear elements, where for surfaces we follow the setup of [52]. An *m*-dimensional manifold *M* is approximated by a triangulated *m*-dimensional domain denoted by *M*_*h*_. The elements of the discretization will be distinguished based on their codimension. We suppose *M*_*h*_ is composed of a collection *T*_*h*_ of *m*-simplices which vertices ***y***_*i*_, *i* = 1, …, *N* lie on *M* and such that

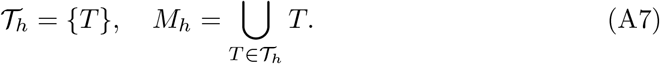

For each *T* ∈ *T*_*h*_ we denote by *h*(*T*) its diameter and define the global *mesh-size* as

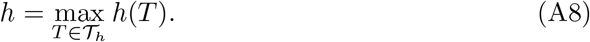

The boundary of every element *T* is composed by *m* + 1 facets of dimension *m* − 1, forming a collection *F* = {*F*}. The discrete boundary of *M*_*h*_ is defined as the union of those facets whose defining points lie on *∂M* and is denoted *∂M*_*h*_.

Throughout this work, we use a first-order fitted finite element method to discretize the PDEs in space [105]. In particular, for surfaces, we employ the surface finite element method (SFEM) as presented in [52]. The corresponding finite element space is defined as

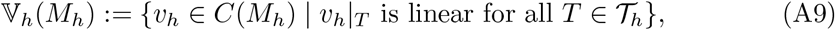

which can be described as the span of *N* piecewise linear continuous basis functions *ϕ*_*i*_, *i* = 1, …, *N* such that *ϕ*_*i*_(***y***_*j*_) = *δ*_*ij*_. Occasionally, we will also need the space

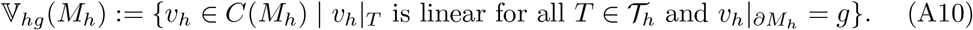

Integration over mesh entities follows the same notation as for the continuous one, where the summation over every element of the collection is implicit.

The evolving manifold *M* (*t*) is discretized using the reference discretization 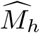 and transporting it using a suitable ALE map **Φ**^*A*^ [58, 104, 106–112]. The reference points 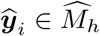 defining the simplices of the reference triangulation *T*_*h*_ are evolved as 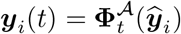.

For the discretization of the function spaces, we start with the finite element space defined in (A9), then transport the basis functions 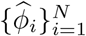 of the reference mesh using the flow map 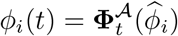 and define the evolved finite element space as their span

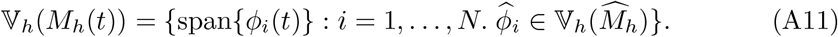

The space V_*h*0_(*M*_*h*_(*t*)) is defined analogously. It follows from the definitions that 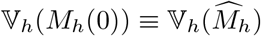. Discrete material velocities are defined by

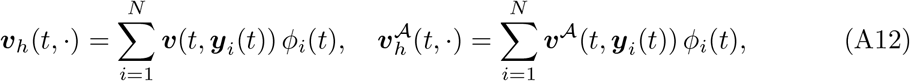

with corresponding derivatives

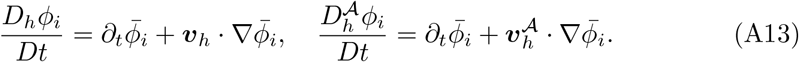

Using an ALE map, which describes the motion of the computational domain, it holds that 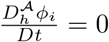.

In this work, time-dependent PDEs are discretized in a collection of time points {*t*_0_, …, *t*_*N*_} . We denote quantities on discrete time points by a superscript, where (·)^*n*^ refers to the current time step and (·)^*n*+1^ refers to the next time step. The time step size is defined as *τ*^*n*+1^ = *t*_*n*+1_ − *t*_*n*_ and for the sake of simplicity we restrict to a constant time step size *τ* ; every one-step method is amenable to adaptive time stepping. We highlight that this time discretization, which is piecewise constant in time, applies to geometric entities as well, for which we write 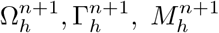, etc.

## Appendix B Numerical algorithms

### B.1 Gradient flow with spatially non-uniform spontaneous curvature

We start from the Helfrich energy functional, which reads

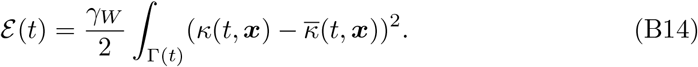

We highlight that here, in contrast to past work, 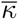 is not assumed constant 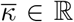, but a function of both space and time 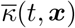. Its variation, found by taking the total ALE time derivative [113, Thm. 2.10.1], can be expressed as follows

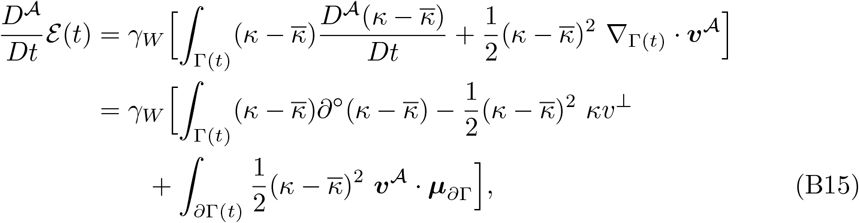

where ***µ***_*∂*Γ_ is the co-normal to the surface boundary. Knowing from the analysis of hypersurfaces [102] that the following holds

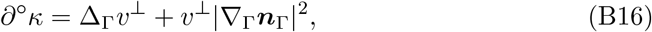

we can plug B16 into the original expression and use integration by parts twice to obtain

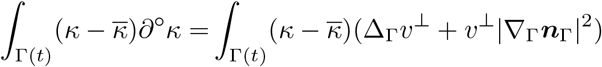

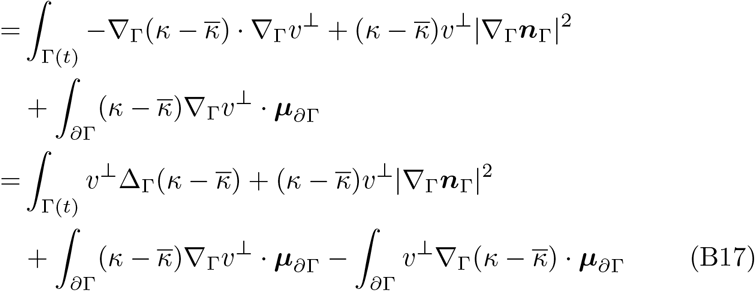

Plugging (B17) back into (B15) yields

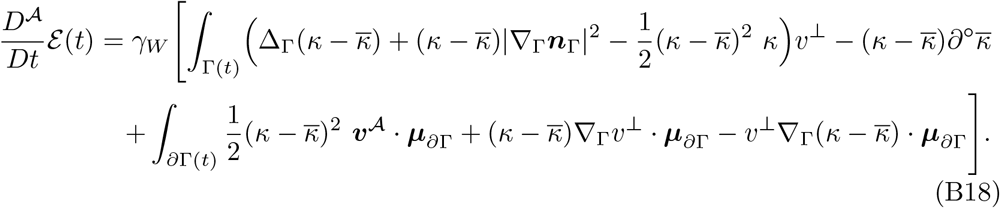

Assuming a fixed boundary, i.e. ***v***^*A*^|_*∂*Γ_ = 0, together with either

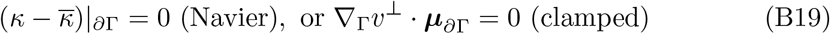

boundary conditions, we can get rid of the whole last line in (B18), ending up with

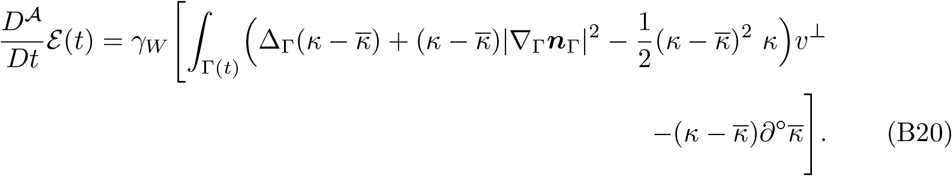

The first line in Equation (B20) constitutes the expression for the Helfrich flow when 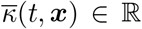. To preserve compatibility while remaining in the larger heterogeneous setting, we then further assume that 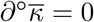. This also implies that

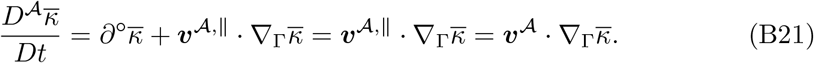

The gradient flow thus has velocity

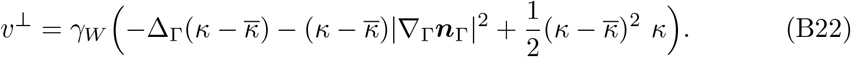

To complete the system we recall the evolution law for the mean curvature [102]

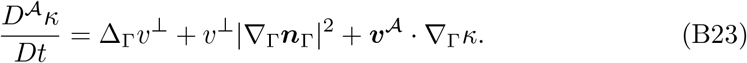

The overall system reads

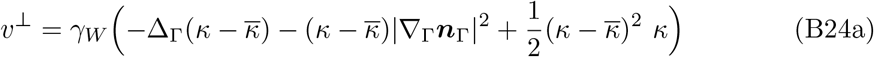

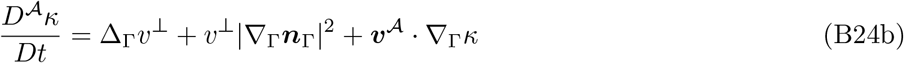

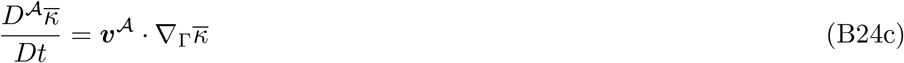

or, equivalently,

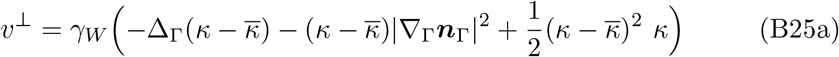

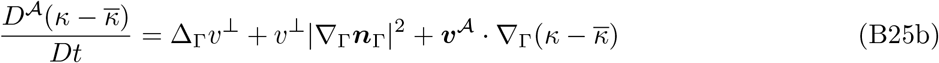

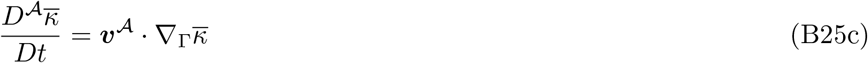

Inspired by the work of [48], we derive the corresponding weak formulation by testing with suitable test functions (*φ, χ, ξ*), respectively. A crucial step in the process is the equality

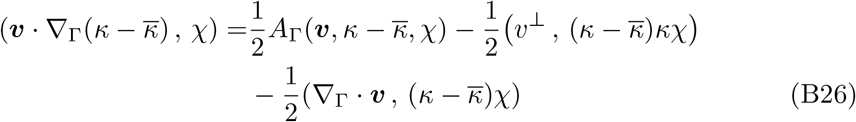

where

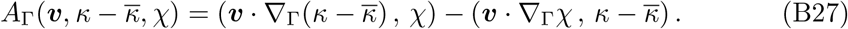

The final weak form reads: find 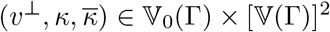 such that

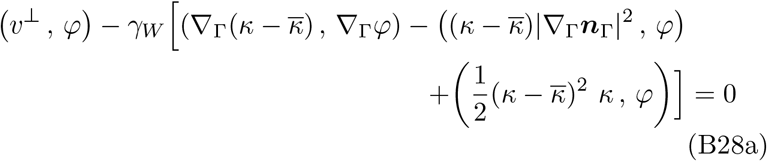

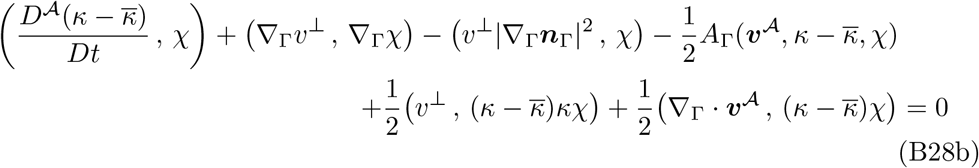

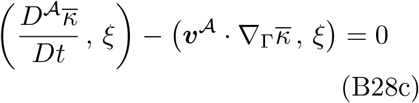

for all (φ, *χ, ξ*) ∈ V_0_(Γ) *×* [V(Γ)]^2^. As in [48], the system (B28) has to be complemented with a system describing the relationship between *v*^⊥^ and ***v***^*A*^. Since we decided to adopt the two-stage strategy of [48], we postpone that second part to Section B.2.

The discretization proceeds as in [48], with the only difference being the additional variable 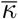, which is now not a constant but evolving with the law in (B21). Supposing the entire boundary 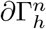 is clamped, the fully discrete scheme reads: find 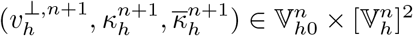 such that

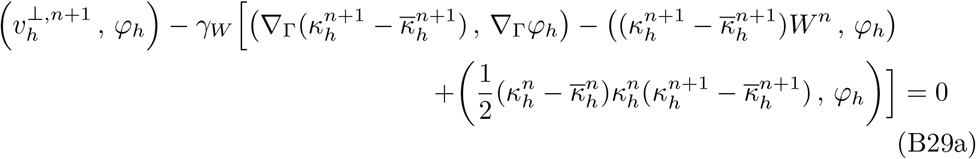

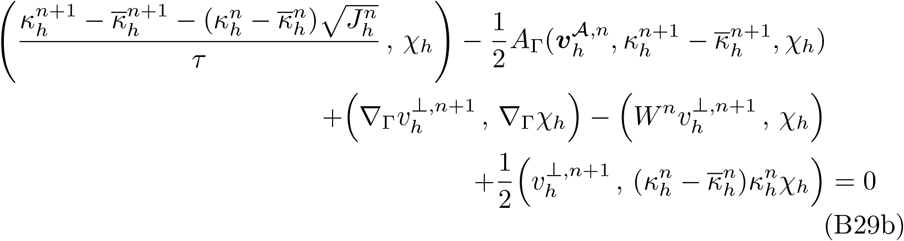

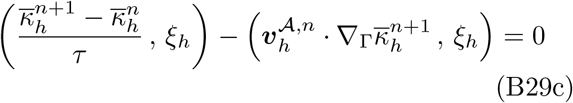

for all 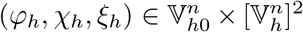, where 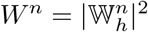 is the square Frobenius norm of the discrete extended Weingarten map 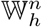 defined on 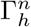. One also needs the discrete ALE velocity 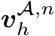 (see Section B.2) between the surfaces 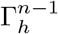 and 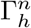 and the change of measure 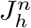 between 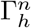 and 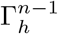 (see [48]).

Using similar arguments as in [48] we can deduce the stability estimate

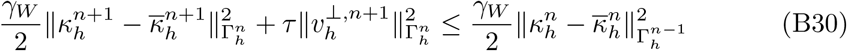

independently of the choice of 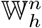. As used by the authors in [48], the initial mean curvature is found by solving the following problem: find 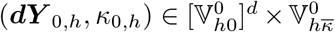 such that

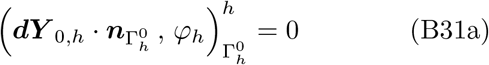

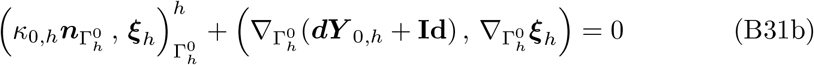

for all 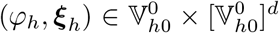. If a stationary solution is needed for a general mesh, we impose 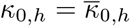.

Equation (B29) constitutes our main force balance equation, as detailed in Section 3.1. The original algorithm of [48] already had the following advantageous properties:

- It allows for both 2D and 3D simulations in a unified setting.
- In its two-stage version a two-scalars-only system is solved in the first step. Moreover, the second stage introduces an independent mesh redistribution which allows for freedom in the choice of the specific redistribution technique (see Section B.2).
- It accounts for surfaces with boundaries with the flexibility of choosing between clamped and Navier boundary conditions.
- It respects, at the discrete level, the energy decaying property typical of the Helfrich flow.

We made the following additions:

- The spontaneous curvature 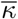 is no longer a constant, but a function 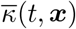 for which it holds that 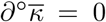. The latter assumption attaches to every material point a characteristic spontaneous curvature. The result is an algorithm allowing for a space-time varying spontaneous curvature that preserves the energy decaying behavior of the continuous equations at the discrete level. In particular, the stability estimate in Equation (B30) introduced in [48] still holds.
- A mean curvature driven gradient flow is added to allow for richer dynamics.
- A multiplier 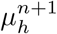 is introduced to allow for volume-preserving flows.

We remark that the new extension preserves the stability estimate in Equation (B30) when Helfrich flow with heterogeneous spontaneous curvature is simulated (i.e. *γ*_*M*_ = 0 and no volume preservation). The final algorithm reads: find 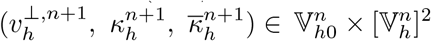 such that

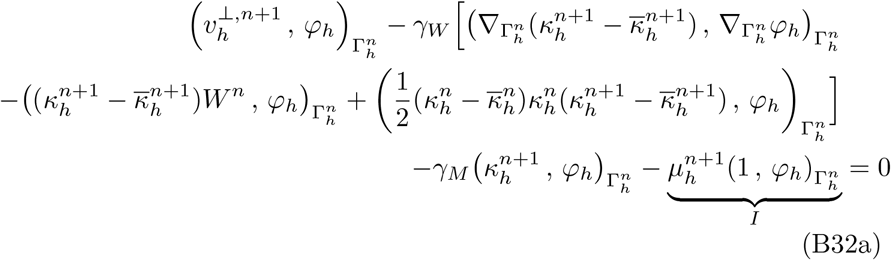

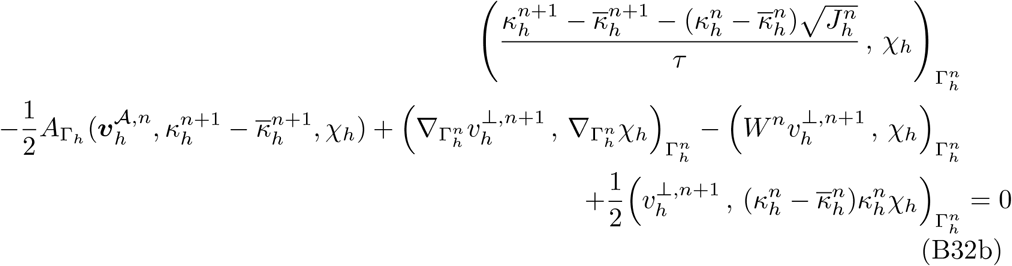

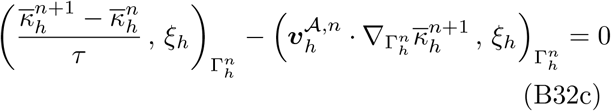

for all 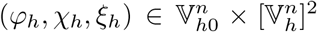. The contribution brought by *γ*_*M*_ incorporates a mean curvature-type flow. Term *I* makes the scheme nonlinear, given that 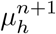 has to preserve the constraint 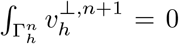, which depends on 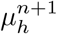 itself. We solved this nonlinear system by taking fixed-point iterations internally until the convergence of both 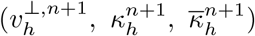 and 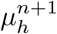.

*Remark 2* The assumption 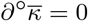 means that 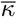 remains constant along the evolution path of every material point.

*Remark 3* In the above we assume that every boundary *∂*Γ_*h*_, even if part of a larger bulk mesh Ω_*h*_, is a fixed boundary with clamped or Navier boundary conditions.

*Remark 4* Notice that the above algorithm is only dependent on the geometric configuration at time *t*_*n*_ and not on the updated geometry at time *t*_*n*+1_. This is a consequence of the specific choice of the discretization, and allows for a decoupling between the geometrical flow solver and the ALE redistribution step.

### B.2 ALE geometric update

Once the force balance has been satisfied (Section 3.1, Section B.1) and 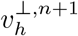 has been computed, the geometrical update has to be performed (Section 3.2). The general idea is to maintain the normal displacement dictated by 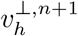, which uniquely prescribes the surface evolution, but redistribute the nodes tangentially to maintain mesh quality. Such an approach, proposed in different forms [59, 63], has proven to be crucial for stable simulations of gradient flow dynamics [49, 51, 114].

Considering a timestep interval [*t*_*n*_, *t*_*n*+1_], we introduce the following quantities:

- The function 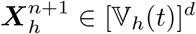 is the discrete analogue to the ALE map 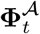, while the function 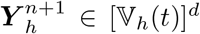 encodes the (total) displacement. Given the nodal basis functions *ϕ*_*i*_(*t*) associated with the evolving mesh points ***y***_*i*_(*t*), they are defined as:

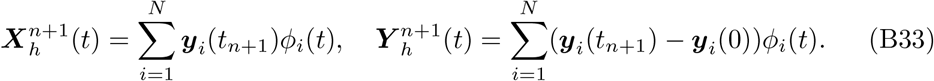
- The discrete displacement between two domains 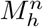 and 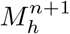 is denoted 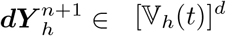, for which one has

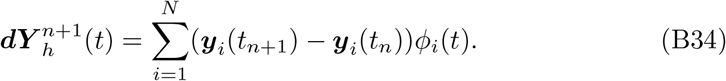

Since we are employing first order schemes, this also defines the piecewise constant discrete velocity 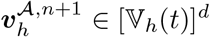 as

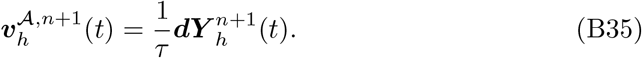

The ALE surface displacement used in this article is taken from [63] and computed as follows. Find 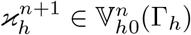 and 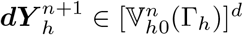 such that

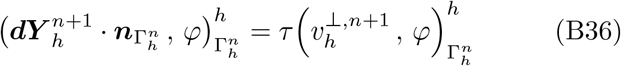

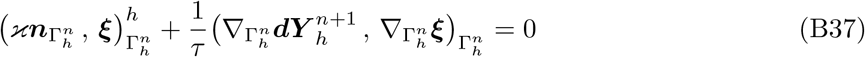

for all 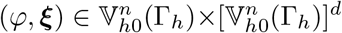. The inner product (·, ·)^*h*^ indicates a mass-lumped inner product.

After the surface has been displaced, a second step might be needed to extend the ALE motion in the bulk. We chose to advect the bulk mesh by solving a harmonic problem. The formulation reads: find 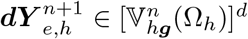

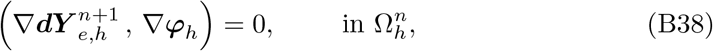

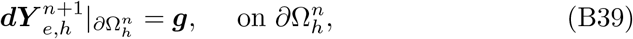

for all 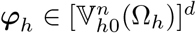, with 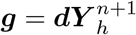.

*Remark 5* As derived in [48], the geometric update described here does not interfere with the energy-decaying property of the Helfrich flow derived in Section B.1. This was not the case for the two-stage procedure proposed in [59], where the tangential motion could potentially destroy the energy-decaying property of the Willmore algorithm in the first stage.

*Remark 6* The material velocity 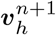 can be computed as 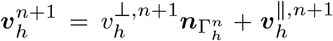 where 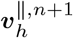 encodes the material tangential motion (usually given by an additional equation such as a tangential fluid equation). Once 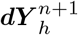 and thus 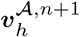 has been computed it can also be useful to define the relative velocity 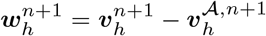, that describes the relation between the material and ALE map.

#### B.3 Advection-diffusion-reaction equations

For the sake of generality here we focus on *moving surfaces with boundary*, for which the equation of a single species reads (see Section 3.3)

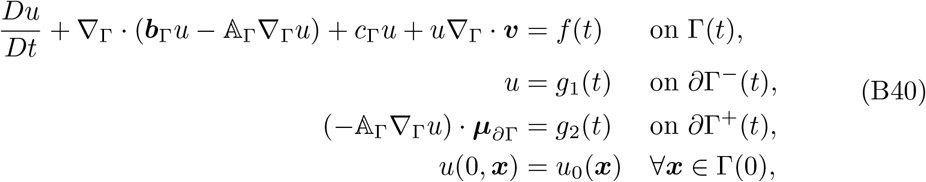

where *∂*Γ^−^(*t*) and *∂*Γ^+^(*t*) are the inflow (***b***_Γ_ · ***µ***_*∂*Γ_ ⩽ 0) and outflow (***b***_Γ_ · ***µ***_*∂*Γ_ *>* 0) boundary, respectively. The semi-discretized in space problem reads: Find *u*_*h*_ ∈ V_*h*_(Γ_*h*_) such that

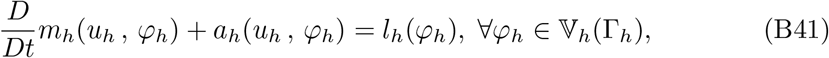

where

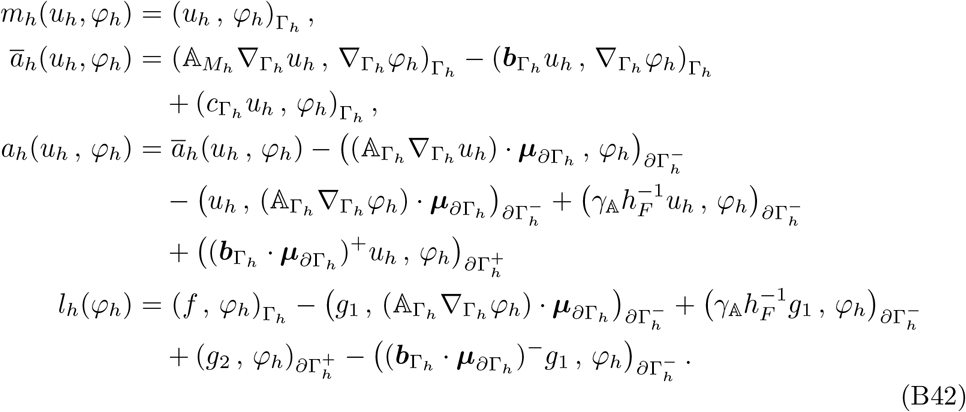

The notation *v*^+^ = max{*v*, 0} and *v*^−^ = min{*v*, 0} has also been used. The case of Robin boundary conditions can readily be adapted. The parameter *γ*_A_ *>* 0 is *Nitsche*’*s penalty parameter* enforcing 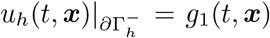 weakly on 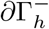 and *h*_*F*_ denotes the facet size.

To obtain the fully discrete version, we use the midpoint time discretization presented in [115], which reads

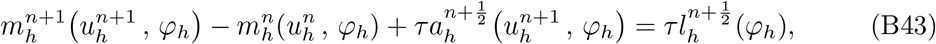

for all Φ_*h*_ ∈ V_*h*_(Γ_*h*_). Such a discretization allows to satisfy the Geometric Conservation Law introduced in [116]. We refer to [112] for a review.

We treated every nonlinearity explicitly by including them in the right-hand side (see Section C), moreover in case the ALE and material maps differ, we compensated with the additional tangential relative velocity 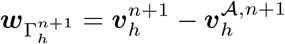.

*Remark 7* Note that the bilinear forms require the updated geometry at times *t*_*n*+1*/*2_ and *t*_*n*+1_.

*Remark 8* Note that the discrete ALE velocity 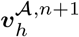 (see (B35)) is piecewise constant in time, and thus independent of the integration domain time for the interval [*t*_*n*_, *t*_*n*+1_].

## Appendix C Numerical testing

We present here convergence tests that we performed to ensure the correctness of our algorithms, divided by application for simplicity. The CIP interior-penalty term is omitted from the displayed ADR bilinear forms for brevity.

### C.1 Application 1

The first step was to verify the convergence properties of our ADR scheme for the case of a moving domain. We tested a coupled surface-bulk equation in the form:

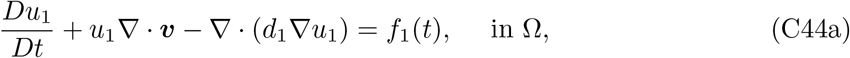

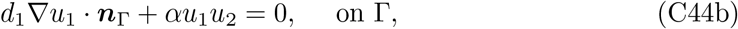

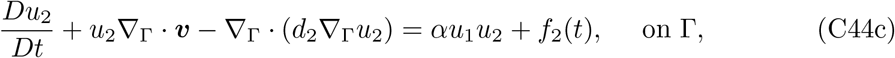

where ***v*** is the material velocity, *α* ∈ R is a coupling parameter and *d*_1_, *d*_2_ are diffusion constants. The domain evolution was described by a map ***x*** = **Φ**(*t*, ***p***) given by

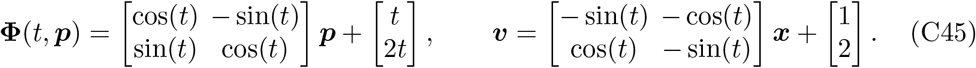

The test is performed on a circle of unit radius and parameters are set as in Table C1.

**Table C1.**

| $t_{end}$ | $\alpha$ | $d_1$ | $d_2$ |
| --- | --- | --- | --- |
| 1 | 2 | 1 | 1 |

Using the right-hand sides *f*_1_, *f*_2_, manufactured solutions were imposed so that *u*_1_ = sin(*t*) cos(*x*_1_*x*_2_) and *u*_2_ = sin(*t*) sin(*x*_1_*x*_2_). Convergence results are presented in Figure C.1a.

Furthermore, we tested a manufactured solution example from a coupled simulation where the domain evolution is unknown. The convergence study was originally presented in [117, 118] and reused in [55, Section 8.1], from which we take the parameters.

The system which is solved is the following [118, p.686]:

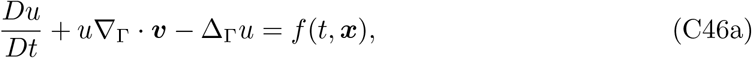

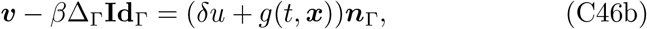

with parameters *β, δ* ∈ ℝ^+^ and ***x*** = (*x*_1_, *x*_2_, *x*_3_). It represents a mean curvature flow driven by a species that diffuses on the surface. Convergence studies are performed choosing *f, g* such that the exact solution for *u* is *u*(*t*, ***x***) = *x*_1_*x*_2_*e*^−6*t*^. The geometry is chosen to be a sphere whose radius evolves following the law

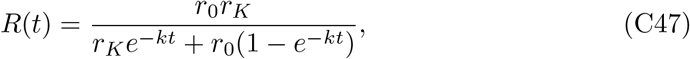

with parameters *r*_0_, *r*_*K*_, *k* ∈ ℝ^+^. The parameters are chosen as in Table C2. The error metric chosen is defined by

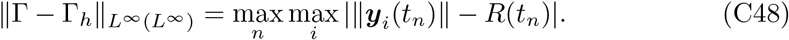

where ***y***_*i*_(*t*_*n*_) is the position of the mesh nodes at time *t*_*n*_. The results are presented in Figure C.1b. The optimal convergence rates are achieved in both space and time for both tests.

**Table C2.**

| $t_{end}$ | $\beta$ | $\delta$ | $r_0$ | $r_K$ | $k$ |
| --- | --- | --- | --- | --- | --- |
| 1 | 1 | 0.4 | 1 | 2 | 0.5 |

Having verified the individual solvers, we present the full coupling algorithm for Application 1 (Section 1.2), following the notation introduced in Sections 3 and 3.4

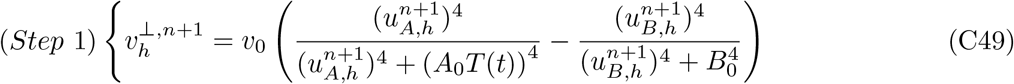

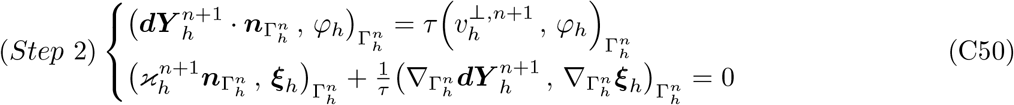

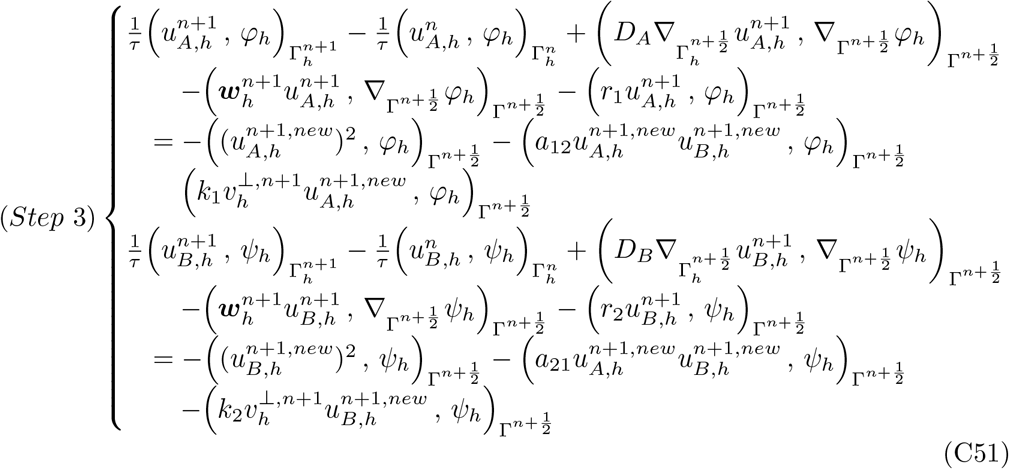

The relative velocity 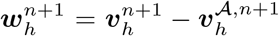 has been introduced to account for the difference between the mesh velocity and the material velocity. As can be observed, the nonlinearities of the ADR system itself are eliminated by evaluating them explicitly, but will still be solved implicitly by the staggering approach. Solving the nonlinearities implicitly internally to the ADR solver is an alternative that we consider in future developments. The initial concentrations are set as 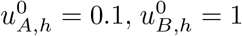. As explained in [12, Supplementary material, “Dependence on initial conditions”], a depletion of the unbranched actin is performed after a short equilibration phase of 10*τ* . This depletion reads

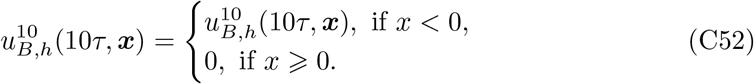

Parameters are taken from Lomakin et al. [12] and are given in Table E3. A first set of preliminary scaling tests is performed and reported in Figure C.7. The results demonstrate linear scaling for this application.

### C.2 Application 2

The technical challenges encountered in Application 2 are the introduction of a surface-bulk coupling and geometric PDEs governing the evolution of an elastic membrane. Since we already tested the surface-bulk convergence properties in the previous section, we focus here on the geometric flow. We start with simple tests of convergence for Willmore flow and we quantify the error with the same metric as in Equation (C48). The first test uses an open Clifford torus with clamped boundary conditions, which is a stationary solution of Willmore flow. After that, we move to the evolution of the unit sphere with *κ* = −2, and evolve it for 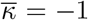 until *t*_*end*_ = 1. The resulting flow can be computed analytically and results in a sphere with evolving radius *R*(*t*) that follows the law

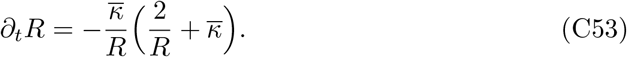

Convergence results are shown in Figure C.2.

To verify the energy decaying and mesh redistribution properties of the newly developed scheme we perform two more tests. The first one is the evolution of a torus with major radius equal to *R* = 2 and minor radius equal to *r* = 1. This geometry does not constitute an energetic minimum and will naturally evolve towards a Clifford torus, which is known to be an energy minimizer with 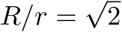 and energy ℰ_*W*_ = 4*π*^2^. To pinpoint a single torus and test the volume-preserving capabilities of the algorithm we decided to also preserve volume. Results for both the scheme in [48] and the novel scheme in Section B.1 are shown in Figure C.3. The second test simulates flow under spontaneous curvature and large deformation regimes. The test is analogous to the one presented in [55], where additional information about the parameters can be found. Two cigar-like surfaces composed of cylinders of height *ℓ* and radius *R* capped with half-spheres are evolved under 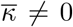. This results in the development of necks and pearls. We test both the case *ℓ/R* = 3 together with 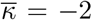 and *ℓ/R* = 5 together with 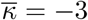. Results are shown in Figure C.4.

As shown in Figure C.2, good convergence properties are attained in both space and time. Moreover, in both Figure C.3 and Figure C.4 the energy-decaying property is satisfied. In particular, the energy for the Clifford torus correctly flattens very close to the expected value and for the pearling case mesh quality is maintained in these extreme necking events. We highlight that results are stable at different mesh resolutions, contrary to previous algorithms, and strictly energy-decaying, a property that was not guaranteed in [55] when mesh redistribution was used.

Having verified the individual solvers, we present the full coupling algorithm for Application 2 (Section 1.3), following the notation introduced in Sections 3 and 3.4

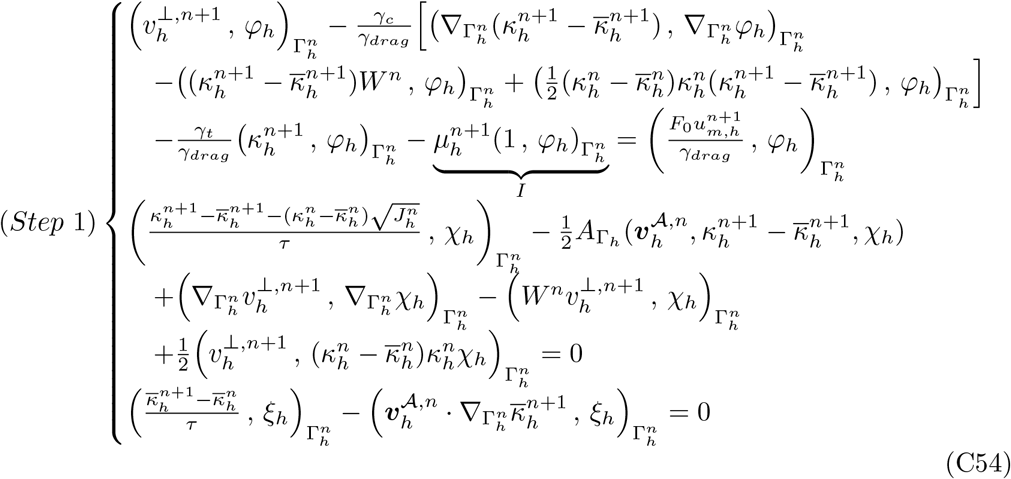

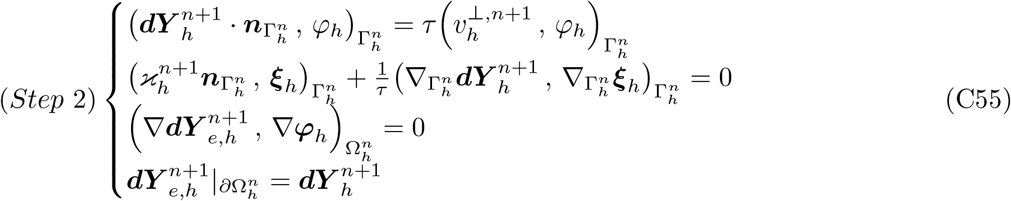

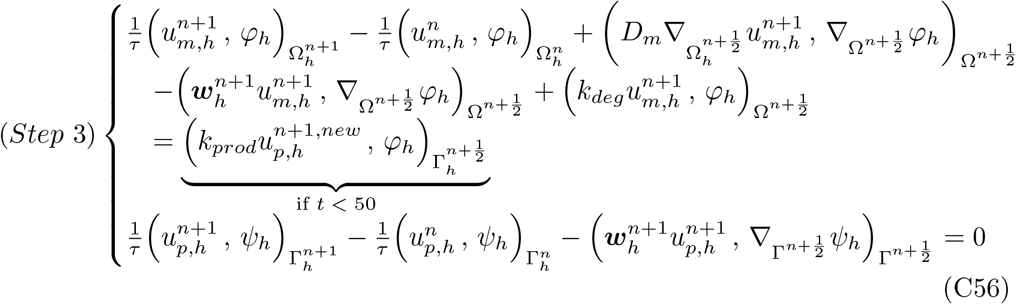

Related parameters can be found in Table E4. A first set of preliminary scaling tests is performed and reported in Figure C.7. The results demonstrate linear scaling in this case. As expected, due to the increased complexity, the per-step time is higher than for Application 1.

#### C.3 Application 3

Application 3 tests the capabilities of our framework by using realistic geometries, where the use of our novel algorithm (Section B.1) is required to guarantee an initial energetic ground state. In the previous sections we demonstrated the accuracy of the surface-bulk solver (Figure C.1a) and the energy-decaying property of the novel elastic algorithm (Figure C.4). What remains to be tested is its applicability to realistic meshes.

We start by running the algorithm on complex meshes and test if the shapes stay stationary. We tried both a closed and an open realistic geometry (Figure C.5a) under different timesteps. Results are shown in Figure C.5. The simulations are very stable and the geometries remain stationary up to machine precision across a wide range of timesteps, including coarse ones. We then tested the energy-decaying properties of the extension in Section B.1 on realistic geometries subjected to external forcing. An open geometry was subjected to a pulling force along the *z*-axis. The force was modeled through an external right-hand side with the form

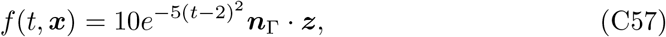

where ***z*** = (0, 0, *x*_3_). The overall simulation was run until *t*_*end*_ = 10 with different timesteps. Results are presented in Figure C.6. One can notice how, under external forcing, the simulations remain stable for a varied set of timesteps, with the Helfrich energy correctly flattening and stabilizing on a certain level once the forcing is no longer active. We remark that the Helfrich flow with 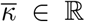 is reparametrization invariant and in general we do not expect the surface to return to the exact zero-energy ground state, since the Helfrich potential ignores stretching phenomena and does not retain a memory of an initial configuration as is the case for other elastic materials. Having assumed 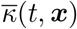 to coevolve with the membrane (i.e., 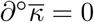), the flow is no longer reparametrization invariant. We thus expect the energy and shape to return to the same initial configuration. Progressive timestep refinement clearly shows convergence towards the initial equilibrium for both energy and relative area change (Figure C.6). We attribute the remaining discrepancies to mesh discretization error.

Having verified the individual solvers, we present the full coupling algorithm for Application 3 (Section 1.4), following the notation introduced in Sections 3 and 3.4

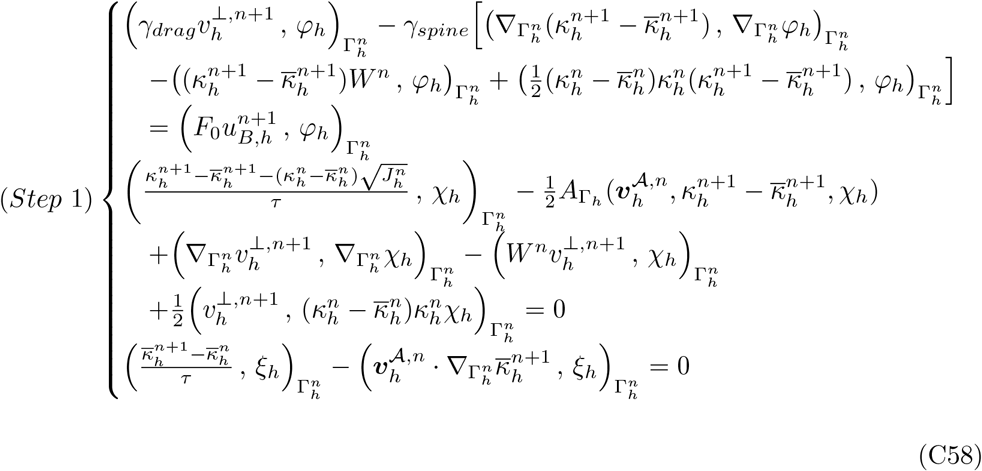

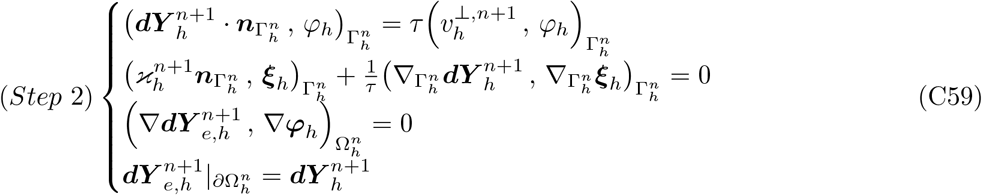

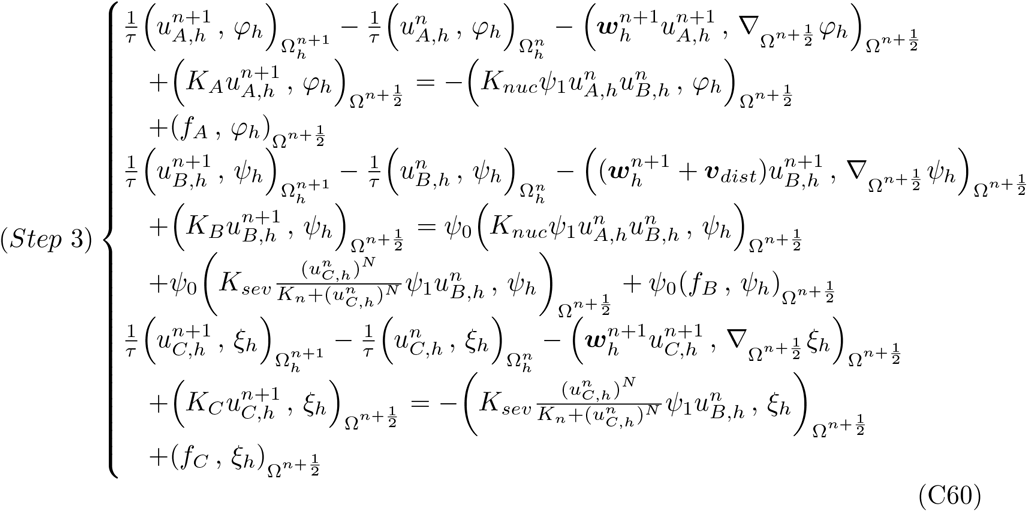

Related parameters can be found in Table E5. Note that for simplicity, contrary to Application 1, the nonlinearities in the ADR system are taken explicitly this time, even considering the staggering scheme. A first set of preliminary scaling tests is performed and reported in Figure C.7. These simulations are more expensive than those in Applications 1 and 2 due to the increased complexity. Nevertheless, linear scaling emerges from our studies. The greater uncertainty in the standard deviation bars is due to the adaptive time stepping procedure, since a discarded timestep size still enters in the count for the total timestep time.

**Fig. C.1:**
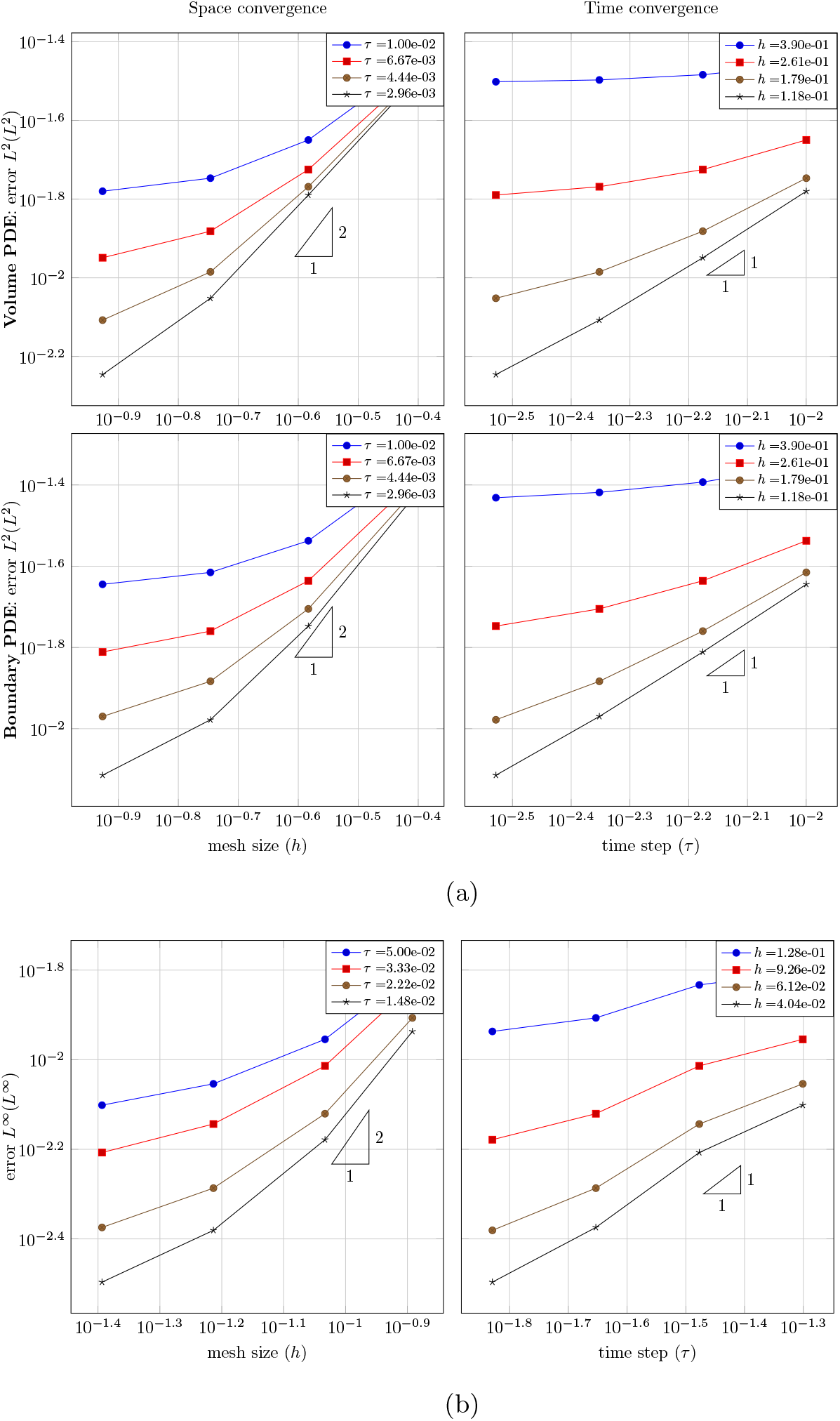
**a)** Space (left) and time (right) convergence for the bulk problem (top) and surface problem (bottom) of the coupled example in Equation (C44). **b)** Space (left) and time (right) convergence of coupled example in Equation (C46).

**Fig. C.2:**
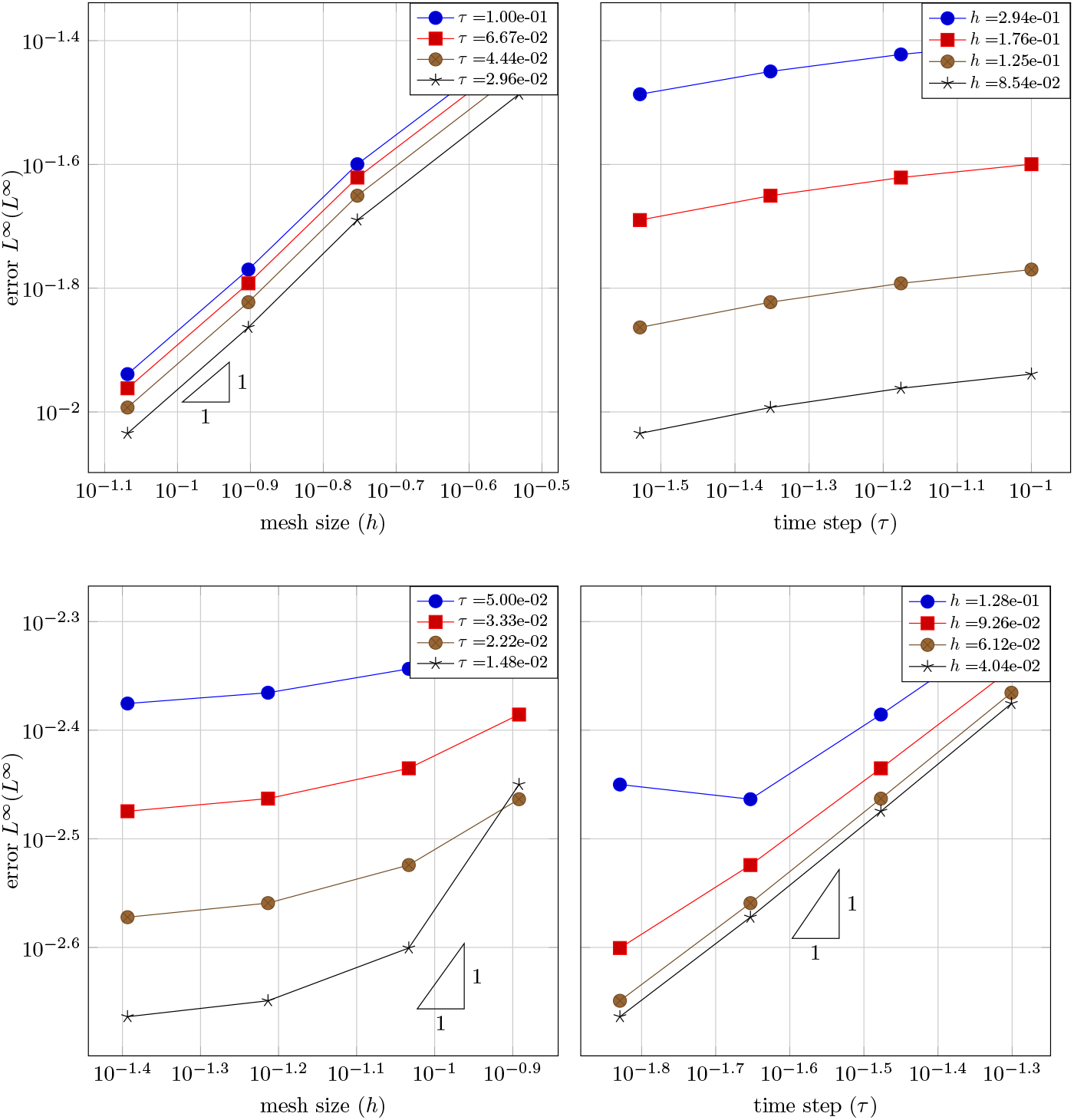
Space (left) and time (right) convergences for manufactured problems for the Helfrich flow algorithm. Top: Half Clifford torus under clamped boundary conditions. Bottom: Sphere under constant spontaneous curvature flow as described in Equation (C53).

**Fig. C.3:**
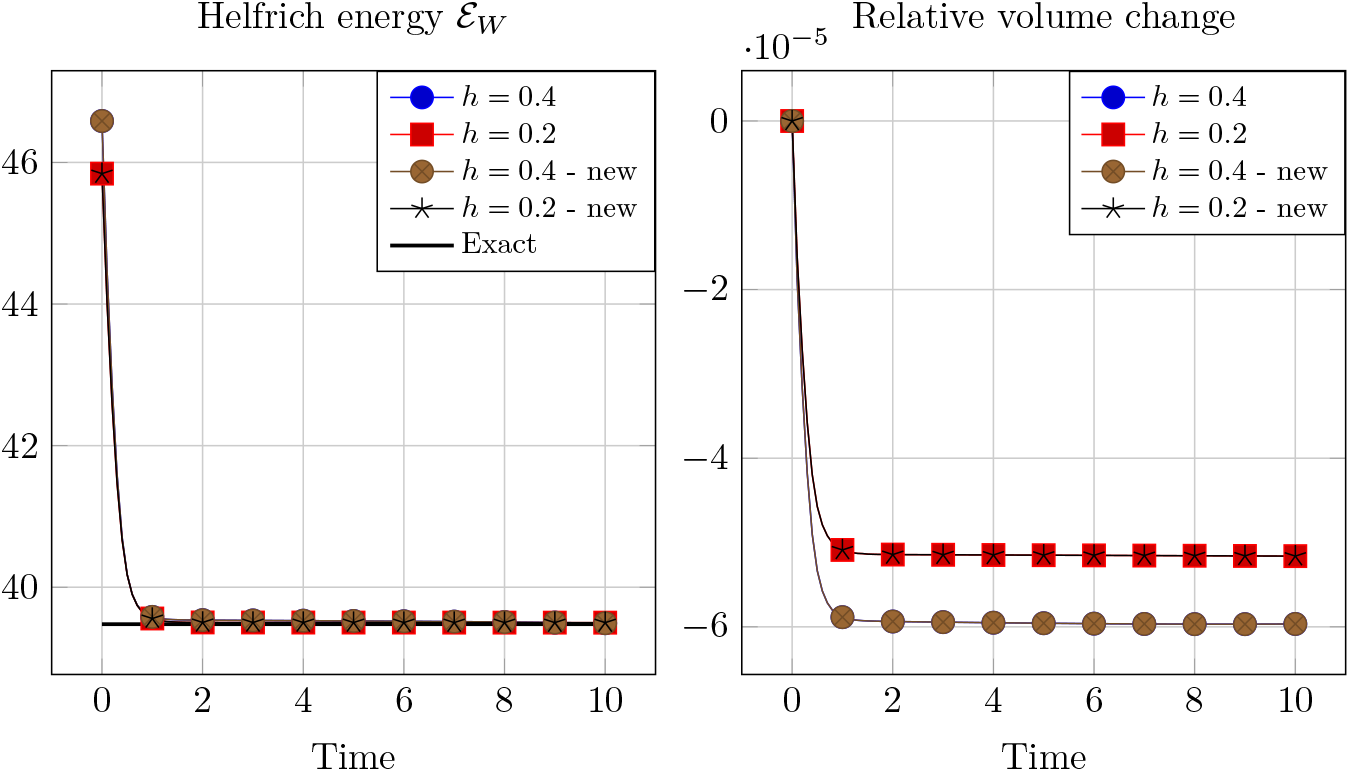
Helfrich energy and volume evolution at different mesh-sizes for a torus with major radius *R* = 2 and *r* = 1. Both the algorithm [48] and the new algorithm in Section B.1 are presented. Exact line in the energy plot is the exact energy for a Clifford torus.

**Fig. C.4:**
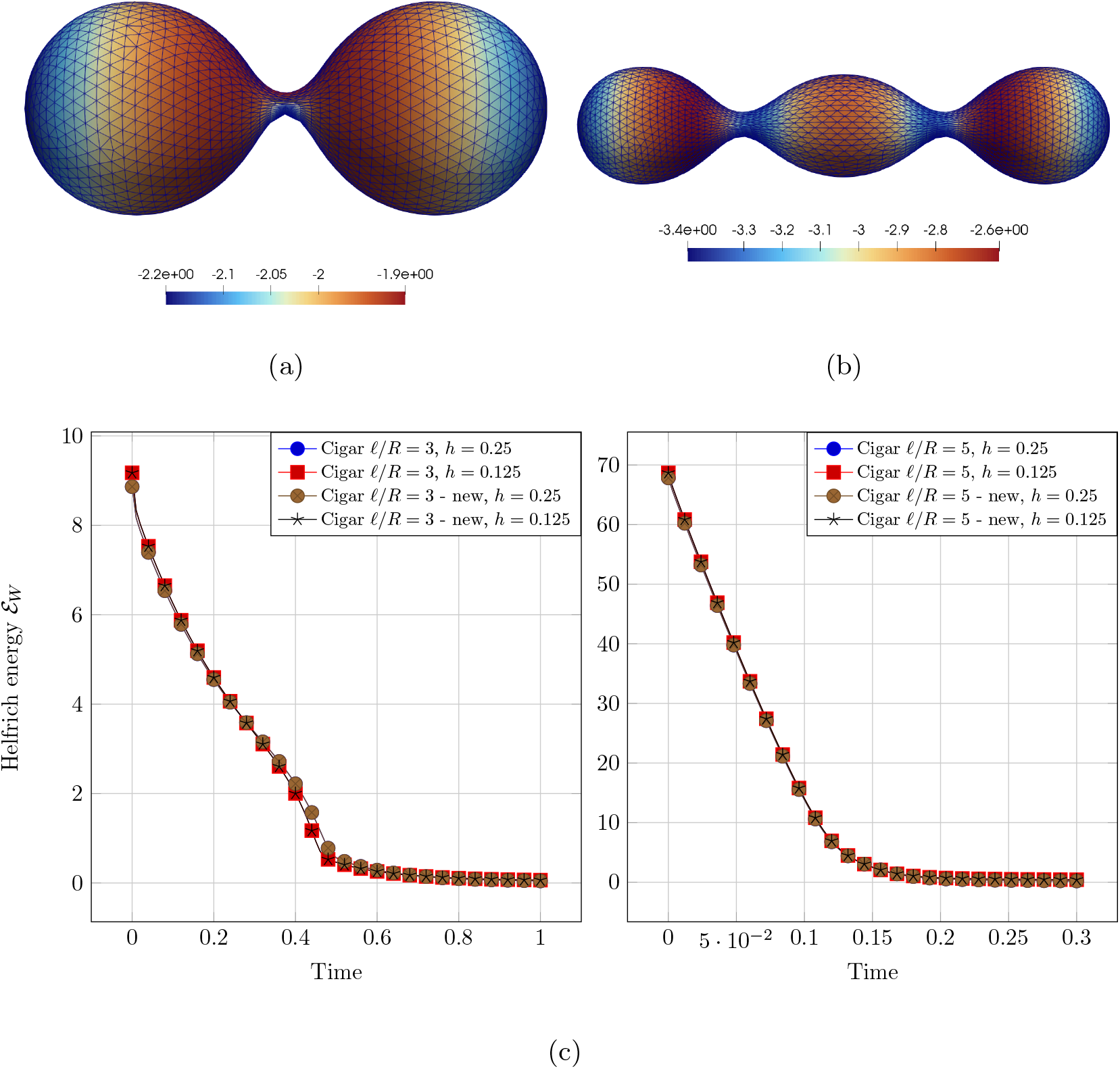
**a)** Final snapshot of Helfrich flow of cigar-like shape with ratio *ℓ/R* = 3 and 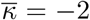. Colorbar shows the mean curvature *κ*. **b)** Final snapshot of Helfrich flow of cigar-like shape with ratio *ℓ/R* = 5 and 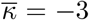. Colorbar shows the mean curvature *κ*. **c)** Helfrich energy evolution for the cigar-shaped examples under different mesh sizes using both the algorithm as in [48] and the novel adaptation of Section B.1. We remark that for the new schemes the spontaneous curvature, instead of being a fixed parameter of the simulation, is set at the beginning only and then co-evolves with the mean curvature. Nevertheless, the results are basically identical between the two.

**Fig. C.5:**
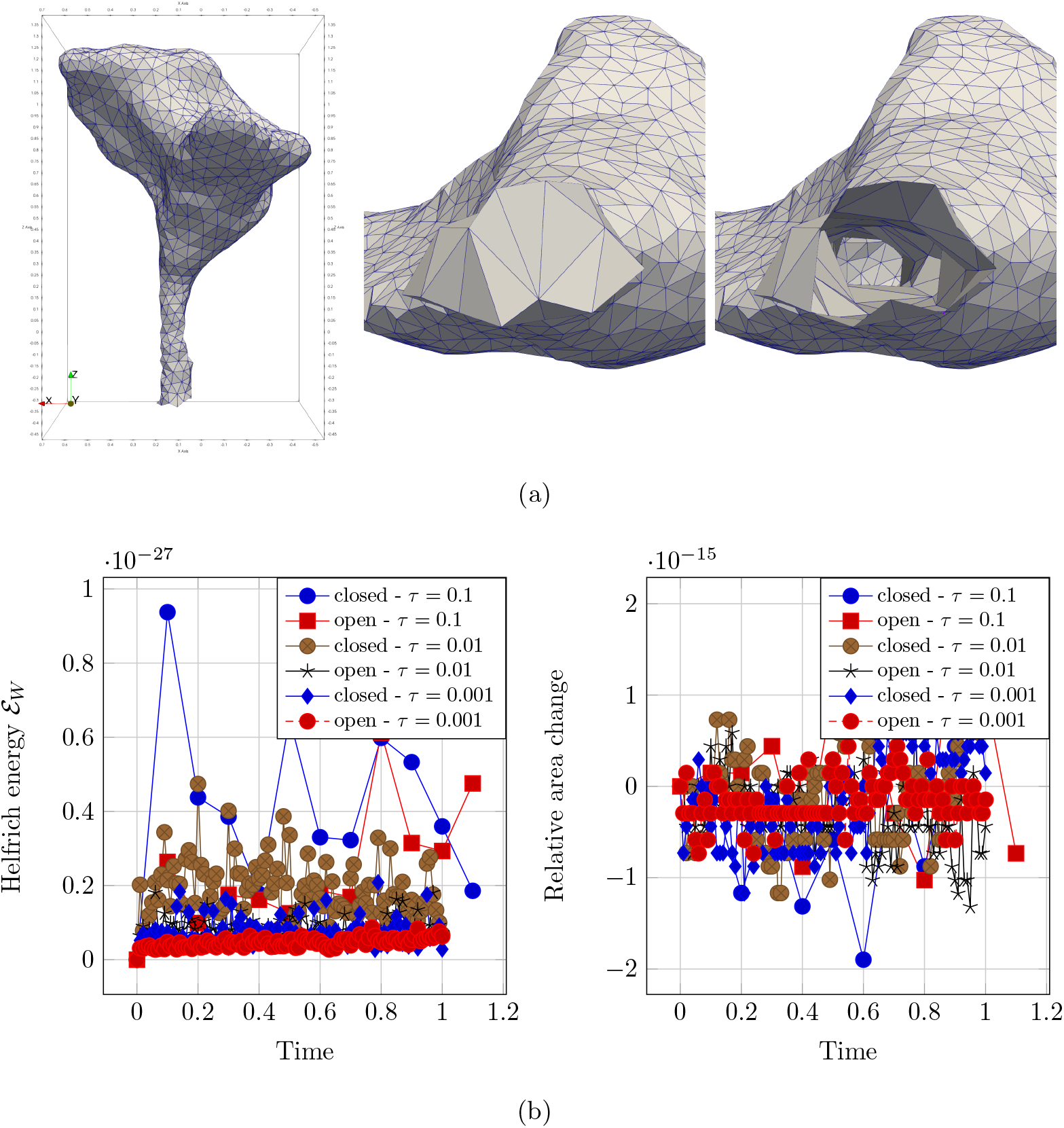
**a)** Realistic meshes for stationary test. Dendritic spine geometry (left, [119]) with closed neck (center, view from the bottom) and open neck (right, view from bottom). **b)** Stability tests for general meshes under Helfrich flow and no external forcing. Tests are presented for a closed and an open surface using different timesteps. Stability is maintained at machine precision.

**Fig. C.6:**
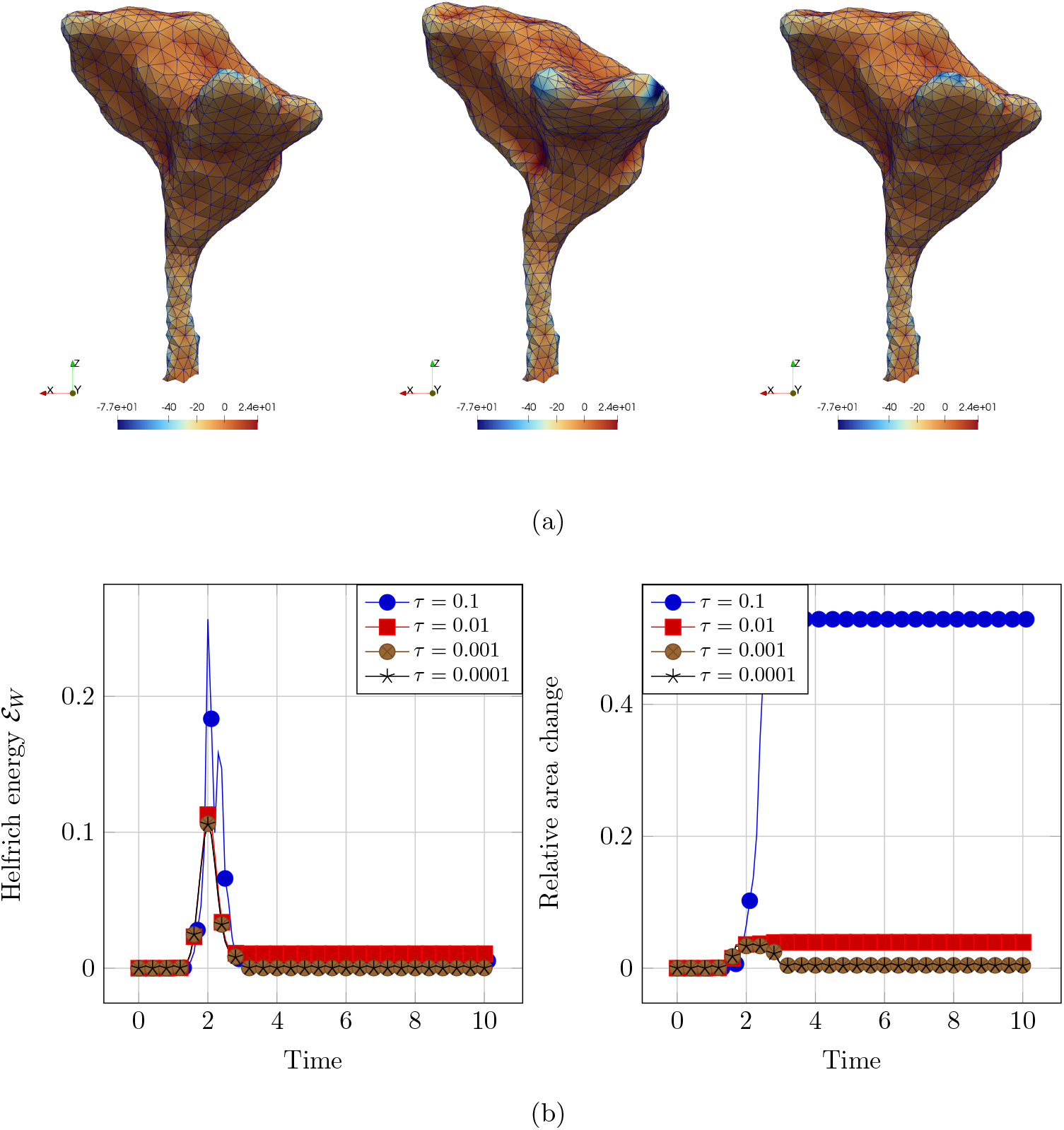
**a)** Geometry for the pull test of Helfrich flow under external forcing at various times: *t* = 0 (left), *t* = 2 (center) and *t* = 10 (right). Colorbar shows the value of the mean curvature. **b)** Stability tests for general meshes under Helfrich flow and added external forcing. Tests are presented for the open surface using different timesteps. Stability is maintained for all timesteps and convergence towards a unique stable configuration is recovered with timestep refinement. Final value of the energy for *τ* = 0.0001 is ℰ_*W*_ ≈ 10^−4^.

**Fig. C.7:**
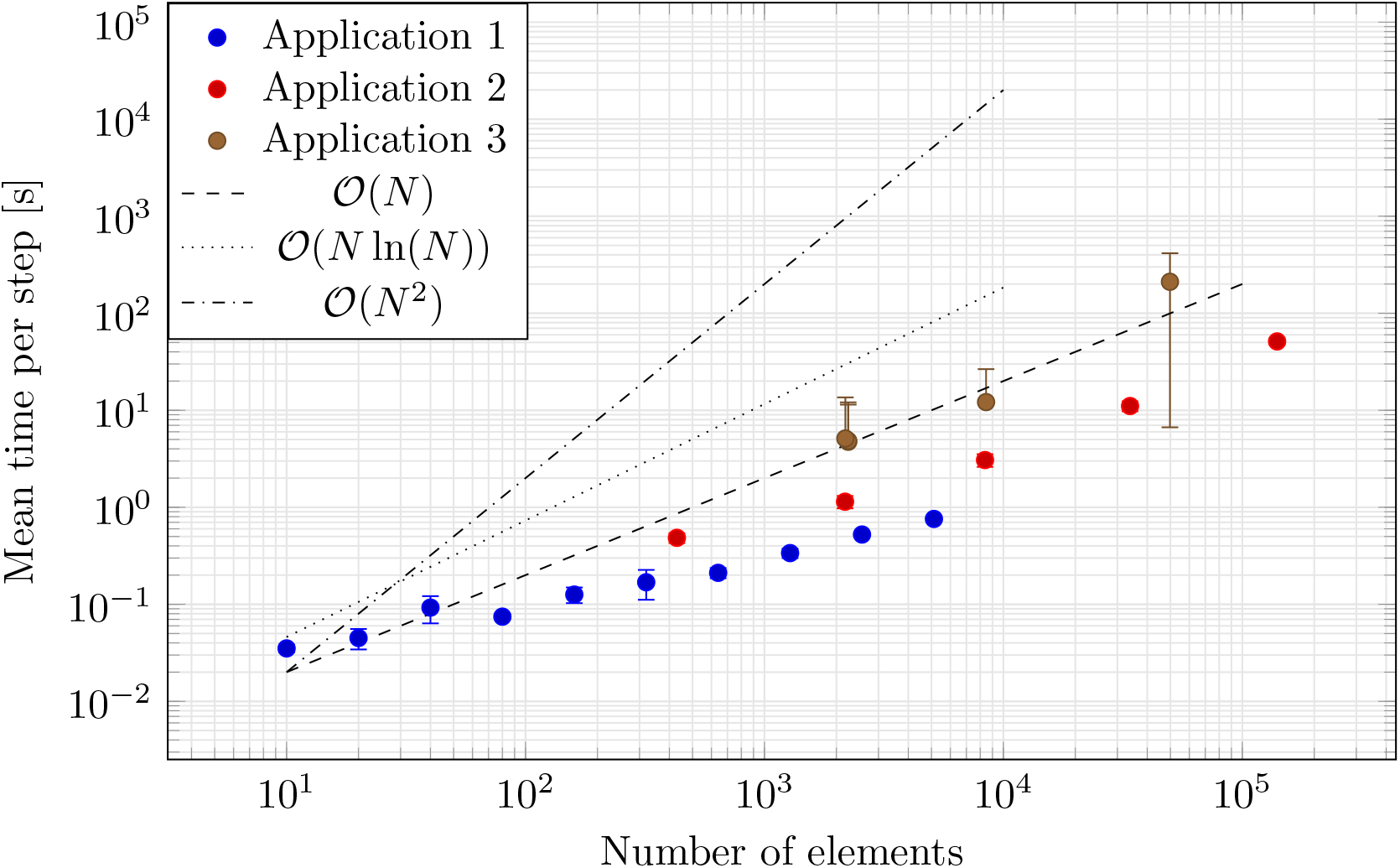
Scaling studies for the tests in Section 1.2, 1.3, 1.4.

## Appendix D Additional supplementary figures

**Fig. D.1:**
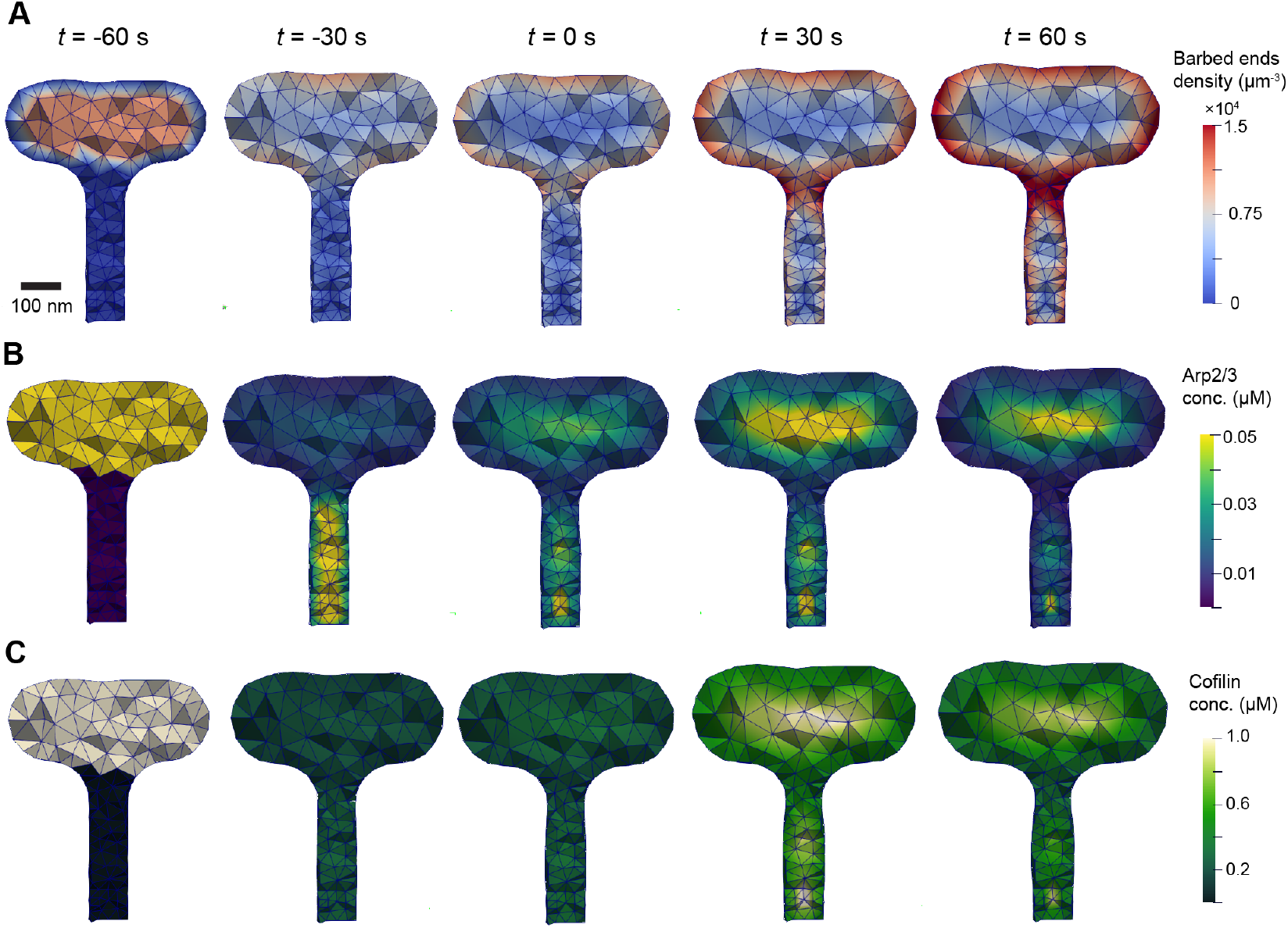
Simulations of dendritic spine plasticity using an idealized spine geometry. The three main biochemical species are visualized within the bulk cytosolic domain, including barbed ends (A), Arp2/3 (B), and cofilin (C). The idealized geometry was designed based on simulations conducted in [81]. It is obtained as a solid of revolution of the profile `wp = occ.WorkPlane().Line(0.05).Rotate(90).Line(0.3).Arc(0.1, -90).Arc(0.1, 180).Line(0.15).Rotate(90).Line(0.6)` following Netgen Python wrapper syntax around the Open Cascade Technology (OCCT) geometry kernel [120].

**Fig. D.2:**
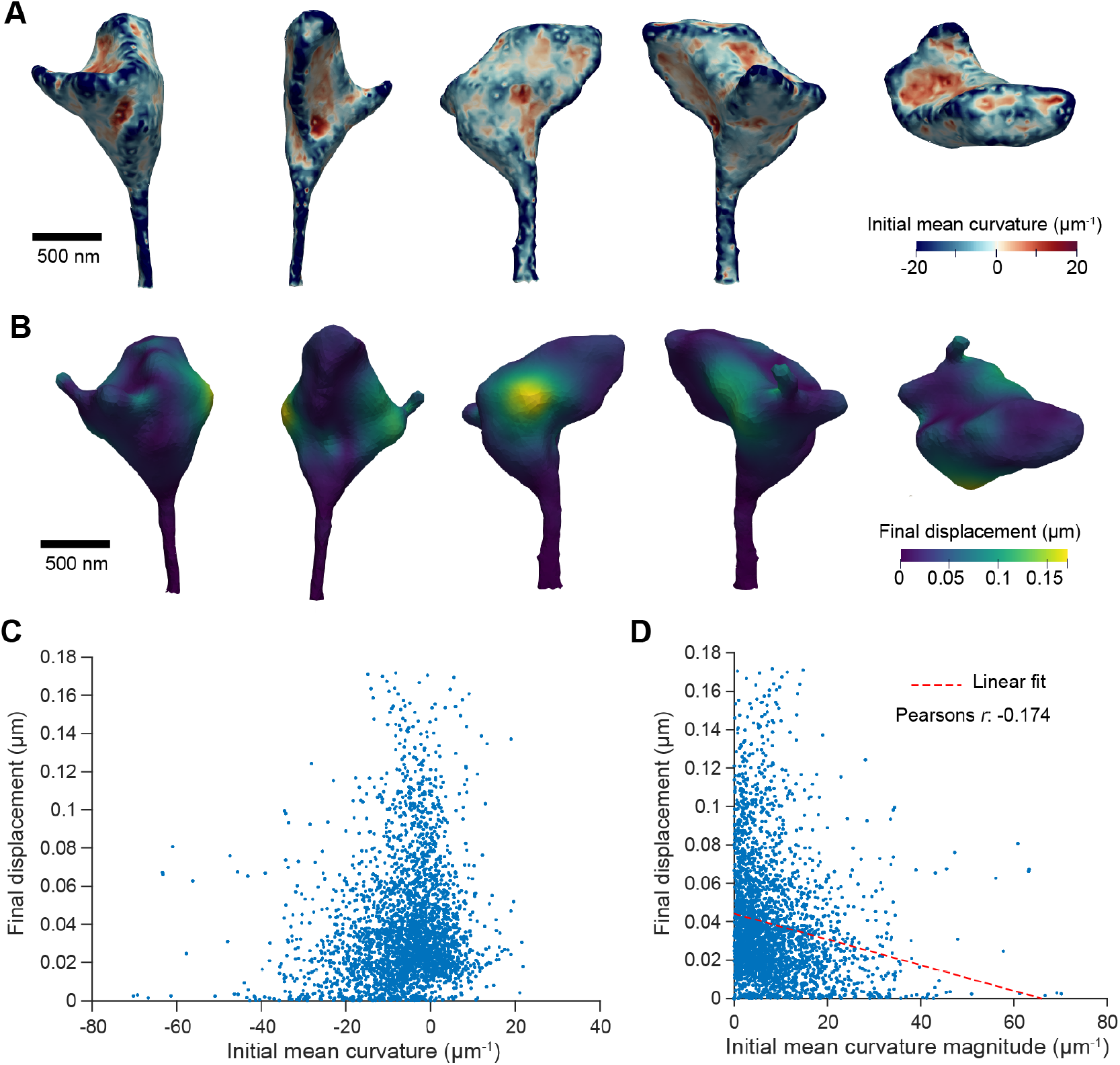
Correlation between normal displacement and initial membrane curvature in spine simulations. A-B) Mean curvature in the initial geometry (A) and final displacement (B) plotted over the spine geometry from different perspectives. C) Final displacement plotted versus the signed mean curvature for each node on the spine plasma membrane. D) Final displacement plotted versus the absolute value of the mean curvature, with an associated Pearson’s correlation coefficient reported. The significant negative correlation coefficient had an associated *p*-value of less than 10^−100^.

## Appendix E Tables of model parameters

**Table E3:** Nondimensional parameters associated with Lomakin et al. model [12].

|  | Value | Description | Source(s)/Notes |
| --- | --- | --- | --- |
| $R_{\text{cell}}$ | 10.0 | Initial cell radius | [12] (assuming 10 $\mu\text{m}$, typical cell radius) |
| $D_A$ | 0.1 | Diffusion coefficient for $u_A$ | [12] (assuming 0.001 $\mu\text{m}^2/\text{s}$ based on [121]) |
| $D_B$ | 0.1 | Diffusion coefficient for $u_B$ | [12] (assuming 0.001 $\mu\text{m}^2/\text{s}$ based on in vitro studies [122, 123]) |
| $r_1$ | 1.0 | Autocatalytic growth coefficient for $u_A$ | Assumed to set time-scale, estimated from data [12, 124] |
| $r_2$ | 1.0 | Autocatalytic growth coefficient for $u_B$ | Estimated to be same order as $r_1$ [12] |
| $k_1$ | 0.4 | Outward flow growth rate for $u_A$ | [12] |
| $k_2$ | 0.4 | Inward flow growth rate for $u_B$ | [12] |
| $a_{12}$ | 0-3 | Effective actin bundling rate | Estimated as order unity in [12] |
| $a_{21}$ | 0-3 | Centripetal flow rate of bundled filaments | Estimated as order unity in [12] |
| $v_0$ | 1.0 | Protrusion/retraction characteristic rate | [12] (assuming characteristic rate of 0.01 $\mu\text{m}/\text{s}$) |
| $A_0$ | 2.2 | Reference concentration of A | Numerical testing |
| $B_0$ | 2.2 | Reference concentration of B | Numerical testing |
| $T_0$ | 1 | Characteristic nondimensional membrane tension | Numerical testing |
| $t_{\text{end}}$ | 100 | Simulation endtime | |
| $\tau$ | $5 \cdot 10^{-2}$ | Simulation timestep | |
| $h$ | 0.63 | Mesh-size | |

**Table E4:**
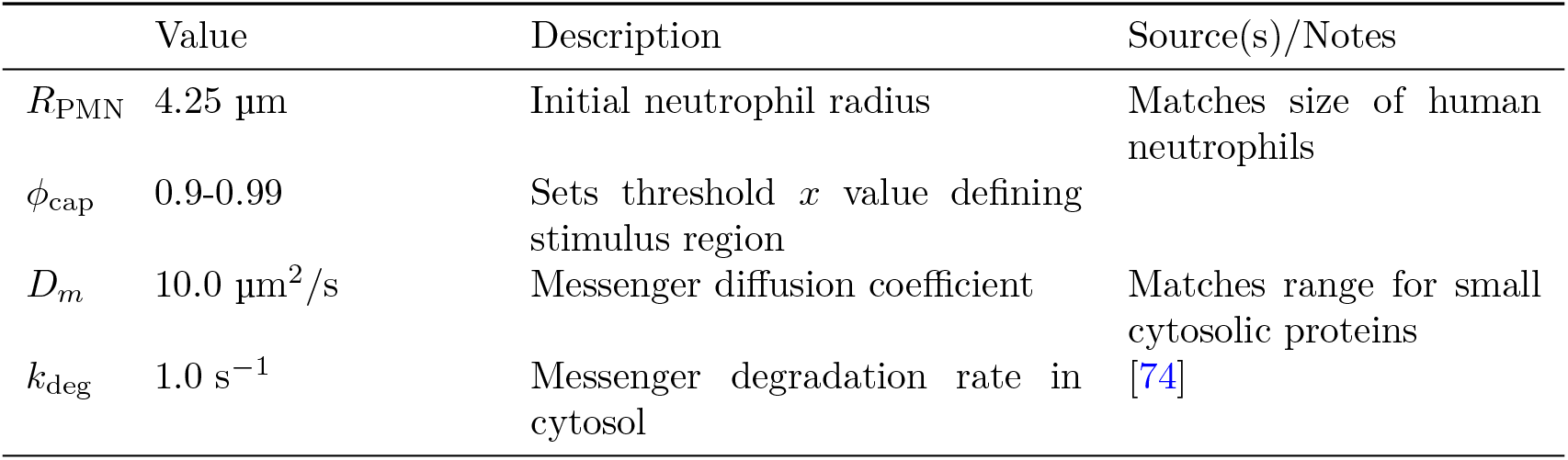
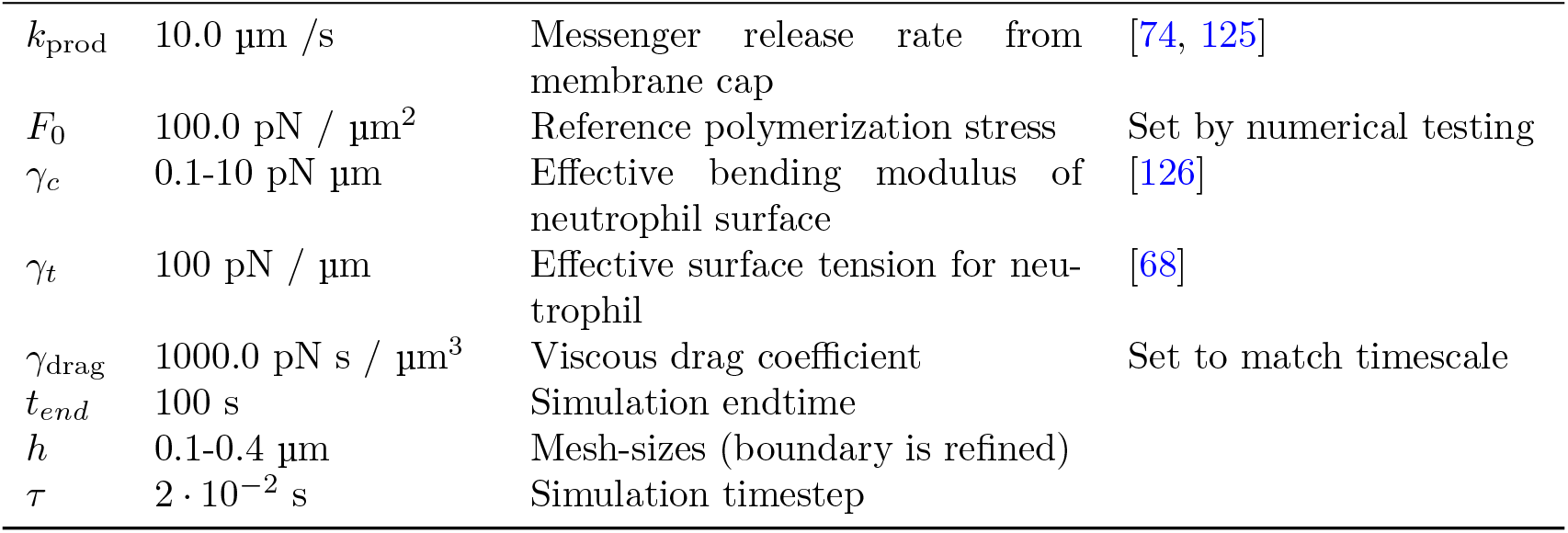
Parameters associated with Herant et al. model.

|  | Value | Description | Source(s)/Notes |
| --- | --- | --- | --- |
| $R_{\text{PMN}}$ | 4.25 $\mu\text{m}$ | Initial neutrophil radius | Matches size of human neutrophils |
| $\phi_{\text{cap}}$ | 0.9-0.99 | Sets threshold $x$ value defining stimulus region | |
| $D_m$ | 10.0 $\mu\text{m}^2/\text{s}$ | Messenger diffusion coefficient | Matches range for small cytosolic proteins |
| $k_{\text{deg}}$ | 1.0 $\text{s}^{-1}$ | Messenger degradation rate in cytosol | [74] |
| $k_{\text{prod}}$ | 10.0 $\mu\text{m} / \text{s}$ | Messenger release rate from membrane cap | [74, 125] |
| $F_0$ | 100.0 pN / $\mu\text{m}^2$ | Reference polymerization stress | Set by numerical testing |
| $\gamma_c$ | 0.1-10 pN $\mu\text{m}$ | Effective bending modulus of neutrophil surface | [126] |
| $\gamma_t$ | 100 pN / $\mu\text{m}$ | Effective surface tension for neutrophil | [68] |
| $\gamma_{\text{drag}}$ | 1000.0 pN s / $\mu\text{m}^3$ | Viscous drag coefficient | Set to match timescale |
| $t_{\text{end}}$ | 100 s | Simulation endtime | |
| $h$ | 0.1-0.4 $\mu\text{m}$ | Mesh-sizes (boundary is refined) | |
| $\tau$ | $2 \cdot 10^{-2}$ s | Simulation timestep | |

**Table E5:**
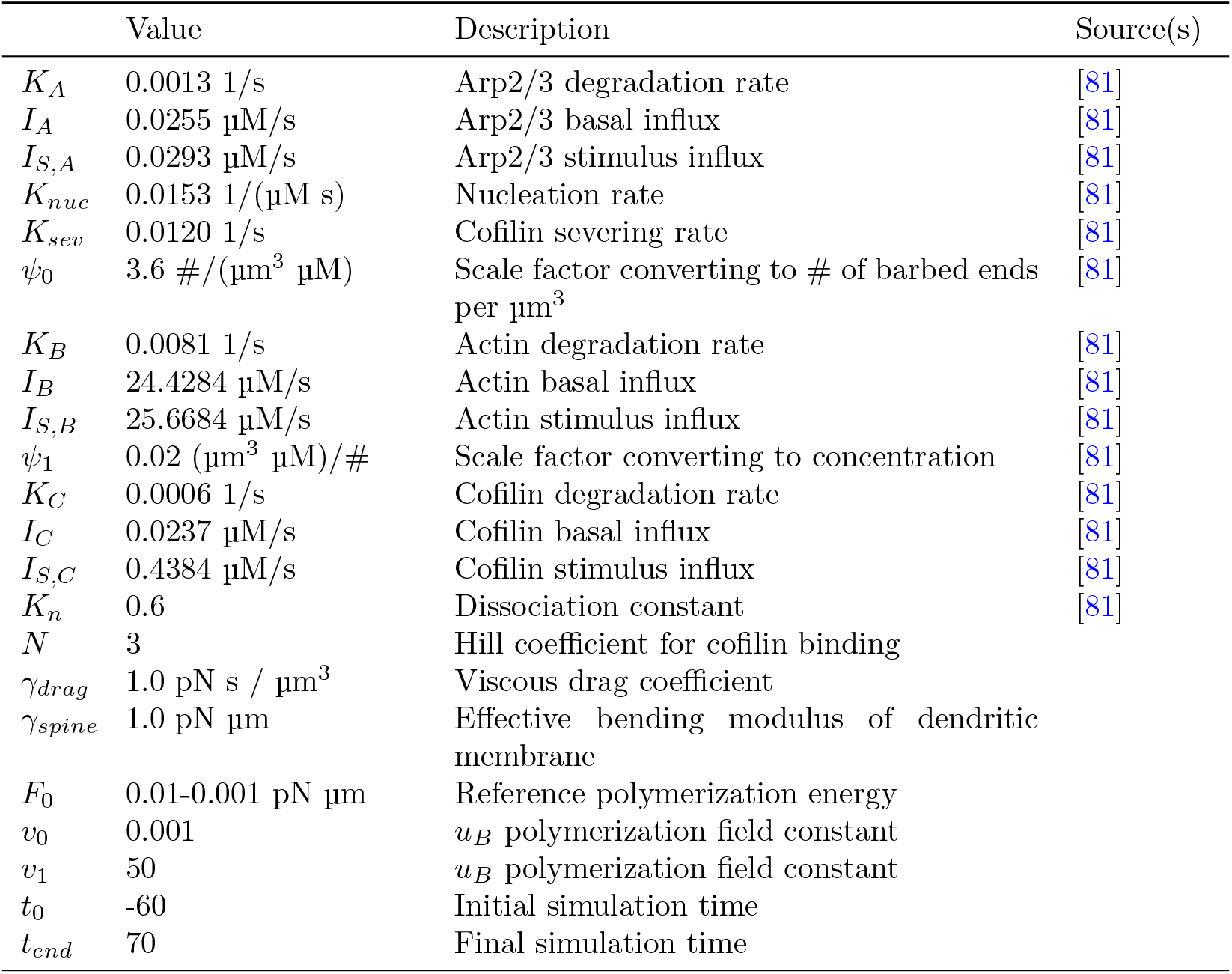
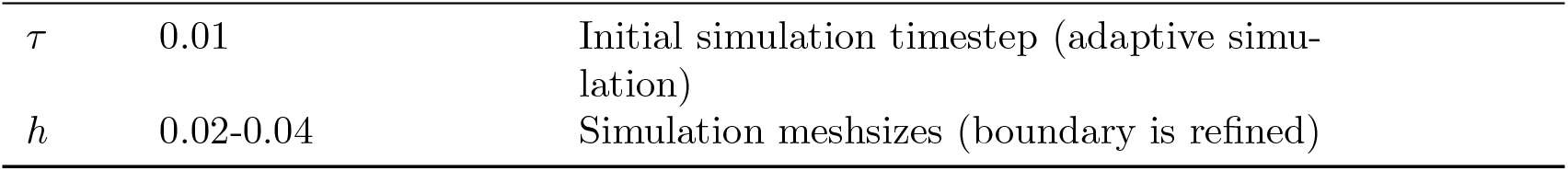
Parameters associated with Bonilla-Quintana et al. model [81].

|  | Value | Description | Source(s) |
| --- | --- | --- | --- |
| $K_A$ | 0.0013 1/s | Arp2/3 degradation rate | [81] |
| $I_A$ | 0.0255 $\mu\text{M}/\text{s}$ | Arp2/3 basal influx | [81] |
| $I_{S,A}$ | 0.0293 $\mu\text{M}/\text{s}$ | Arp2/3 stimulus influx | [81] |
| $K_{nuc}$ | 0.0153 1/($\mu\text{M}$ s) | Nucleation rate | [81] |
| $K_{sev}$ | 0.0120 1/s | Cofilin severing rate | [81] |
| $\psi_0$ | 3.6 #/($\mu\text{m}^3$ $\mu\text{M}$) | Scale factor converting to # of barbed ends per $\mu\text{m}^3$ | [81] |
| $K_B$ | 0.0081 1/s | Actin degradation rate | [81] |
| $I_B$ | 24.4284 $\mu\text{M}/\text{s}$ | Actin basal influx | [81] |
| $I_{S,B}$ | 25.6684 $\mu\text{M}/\text{s}$ | Actin stimulus influx | [81] |
| $\psi_1$ | 0.02 ($\mu\text{m}^3$ $\mu\text{M}$)/# | Scale factor converting to concentration | [81] |
| $K_C$ | 0.0006 1/s | Cofilin degradation rate | [81] |
| $I_C$ | 0.0237 $\mu\text{M}/\text{s}$ | Cofilin basal influx | [81] |
| $I_{S,C}$ | 0.4384 $\mu\text{M}/\text{s}$ | Cofilin stimulus influx | [81] |
| $K_n$ | 0.6 | Dissociation constant | [81] |
| $N$ | 3 | Hill coefficient for cofilin binding | |
| $\gamma_{\text{drag}}$ | 1.0 pN s / $\mu\text{m}^3$ | Viscous drag coefficient | |
| $\gamma_{\text{spine}}$ | 1.0 pN $\mu\text{m}$ | Effective bending modulus of dendritic membrane | |
| $F_0$ | 0.01-0.001 pN $\mu\text{m}$ | Reference polymerization energy | |
| $v_0$ | 0.001 | $u_B$ polymerization field constant | |
| $v_1$ | 50 | $u_B$ polymerization field constant | |
| $t_0$ | -60 | Initial simulation time | |
| $t_{\text{end}}$ | 70 | Final simulation time | |
| $\tau$ | 0.01 | Initial simulation timestep (adaptive simulation) | |
| $h$ | 0.02-0.04 | Simulation meshsizes (boundary is refined) | |

## References

[1] Paluch, E., Heisenberg, C.-P.: Biology and physics of cell shape changes in development. Curr. Biol. 19(17), 790–9 (2009)

[2] McBeath, R., Pirone, D.M., Nelson, C.M., Bhadriraju, K., Chen, C.S.: Cell shape, cytoskeletal tension, and RhoA regulate stem cell lineage commitment. Dev. Cell 6(4), 483–495 (2004)

[3] Luciano, M., Versaevel, M., Vercruysse, E., Procès, A., Kalukula, Y., Remson, A., Deridoux, A., Gabriele, S.: Appreciating the role of cell shape changes in the mechanobiology of epithelial tissues. Biophys. Rev. (Melville) 3(1), 011305 (2022)

[4] Moujaber, O., Stochaj, U.: The cytoskeleton as regulator of cell signaling pathways. Trends Biochem. Sci. 45(2), 96–107 (2020)

[5] Nayak, R.C., Chang, K.-H., Vaitinadin, N.-S., Cancelas, J.A.: Rho GTPases control specific cytoskeleton-dependent functions of hematopoietic stem cells. Immunol. Rev. 256(1), 255–268 (2013)

[6] Sit, S.-T., Manser, E.: Rho GTPases and their role in organizing the actin cytoskeleton. J. Cell Sci. 124(Pt 5), 679–683 (2011)

[7] Argudo, D., Bethel, N.P., Marcoline, F.V., Grabe, M.: Continuum descriptions of membranes and their interaction with proteins: Towards chemically accurate models. Biochim. Biophys. Acta 1858(7 Pt B), 1619–1634 (2016)

[8] Rangamani, P.: The many faces of membrane tension: Challenges across systems and scales. Biochim. Biophys. Acta Biomembr. 1864(7), 183897 (2022)

[9] Boal, D.: Mechanics of the Cell, 2nd edn. Cambridge University Press, Cambridge, England (2012)

[10] Friedl, P., Wolf, K.: Tumour-cell invasion and migration: Diversity and escape mechanisms. Nat Rev Cancer 3(5), 362–374 (2003) 10.1038/nrc1075

[11] Friedl, P., Weigelin, B.: Interstitial leukocyte migration and immune function. Nat Immunol 9(9), 960–969 (2008) 10.1038/ni.f.212

[12] Lomakin, A.J., Lee, K.-C., Han, S.J., Bui, D.A., Davidson, M., Mogilner, A., Danuser, G.: Competition for actin between two distinct F-actin networks defines a bistable switch for cell polarization. Nat Cell Biol 17(11), 1435–1445 (2015) 10.1038/ncb3246

[13] Warner, H., Wilson, B.J., Caswell, P.T.: Control of adhesion and protrusion in cell migration by Rho GTPases. Curr Opin Cell Biol 56, 64–70 (2019) 10.1016/j.ceb.2018.09.003

[14] Othmer, H.G.: Eukaryotic Cell Dynamics from Crawlers to Swimmers. Wiley Interdiscip Rev Comput Mol Sci 9(1), 1376 (2019) 10.1002/wcms.1376

[15] Buttenschön, A., Edelstein-Keshet, L.: Bridging from single to collective cell migration: A review of models and links to experiments. PLOS Computational Biology 16(12), 1008411 (2020) 10.1371/journal.pcbi.1008411

[16] Wang, Y., Li, S., Mokbel, M., May, A.I., Liang, Z., Zeng, Y., Wang, W., Zhang, H., Yu, F., Sporbeck, K., Jiang, L., Aland, S., Agudo-Canalejo, J., Knorr, R.L., Fang, X.: Biomolecular condensates mediate bending and scission of endosome membranes. Nature, 1–7 (2024) 10/g77qdg

[17] Kaplan, C., Kenny, S.J., Chen, X., Schöneberg, J., Sitarska, E., Diz-Mu ñoz, A., Akamatsu, M., Xu, K., Drubin, D.G.: Load adaptation by endocytic actin networks. Mol. Biol. Cell 33(6), 50 (2022)

[18] Watanabe, S., Rost, B.R., Camacho-Pérez, M., Davis, M.W., Söhl-Kielczynski, B., Rosenmund, C., Jorgensen, E.M.: Ultrafast endocytosis at mouse hippocampal synapses. Nature 504(7479), 242–247 (2013)

[19] Zhang, Y., Hoppe, A.D., Swanson, J.A.: Coordination of Fc receptor signaling regulates cellular commitment to phagocytosis. Proc. Natl. Acad. Sci. U. S. A. 107(45), 19332–19337 (2010)

[20] Lee, C.-Y., Herant, M., Heinrich, V.: Target-specific mechanics of phagocytosis: protrusive neutrophil response to zymosan differs from the uptake of antibody-tagged pathogens. J. Cell Sci. 124(Pt 7), 1106–1114 (2011)

[21] Boero, E., Gorham, R.D. Jr, Francis, E.A., Brand, J., Teng, L.H., Doorduijn, D.J., Ruyken, M., Muts, R.M., Lehmann, C., Verschoor, A., Kessel, K.P.M., Heinrich, V., Rooijakkers, S.H.M.: Purified complement C3b triggers phagocytosis and activation of human neutrophils via complement receptor 1. Sci. Rep. 13(1), 274 (2023)

[22] Walbaum, S., Ambrosy, B., Schütz, P., Bachg, A.C., Horsthemke, M., Leusen, J.H.W., Mócsai, A., Hanley, P.J.: Complement receptor 3 mediates both sinking phagocytosis and phagocytic cup formation via distinct mechanisms. J. Biol. Chem. 296(100256), 100256 (2021)

[23] Brangwynne, C.P., Eckmann, C.R., Courson, D.S., Rybarska, A., Hoege, C., Gharakhani, J., Jülicher, F., Hyman, A.A.: Germline P Granules Are Liquid Droplets That Localize by Controlled Dissolution/Condensation. Science 324(5935), 1729–1732 (2009) 10.1126/science.1172046

[24] Banani, S.F., Lee, H.O., Hyman, A.A., Rosen, M.K.: Biomolecular condensates: Organizers of cellular biochemistry. Nat Rev Mol Cell Biol 18(5), 285–298 (2017) 10.1038/nrm.2017.7

[25] Shin, Y., Brangwynne, C.P.: Liquid phase condensation in cell physiology and disease. Science 357(6357), 4382 (2017) 10.1126/science.aaf4382

[26] Beutel, O., Maraspini, R., Pombo-García, K., Martin-Lemaitre, C., Honigmann, A.: Phase Separation of Zonula Occludens Proteins Drives Formation of Tight Junctions. Cell 179(4), 923–93611 (2019) 10.1016/j.cell.2019.10.011

[27] Lee, J.E., Cathey, P.I., Wu, H., Parker, R., Voeltz, G.K.: Endoplasmic reticulum contact sites regulate the dynamics of membraneless organelles. Science 367(6477), 7108 (2020) 10.1126/science.aay7108

[28] Zhao, Y.G., Zhang, H.: Phase Separation in Membrane Biology: The Interplay between Membrane-Bound Organelles and Membraneless Condensates. Dev Cell 55(1), 30–44 (2020) 10.1016/j.devcel.2020.06.033

[29] Alberti, S., Hyman, A.A.: Biomolecular condensates at the nexus of cellular stress, protein aggregation disease and ageing. Nat Rev Mol Cell Biol 22(3), 196–213 (2021) 10.1038/s41580-020-00326-6

[30] Ambroggio, E.E., Costa Navarro, G.S., Pérez Socas, L.B., Bagatolli, L.A., Gamarnik, A.V.: Dengue and Zika virus capsid proteins bind to membranes and self-assemble into liquid droplets with nucleic acids. J Biol Chem 297(3), 101059 (2021) 10.1016/j.jbc.2021.101059

[31] Snead, W.T., Jalihal, A.P., Gerbich, T.M., Seim, I., Hu, Z., Gladfelter, A.S.: Membrane surfaces regulate assembly of ribonucleoprotein condensates. Nat Cell Biol 24(4), 461–470 (2022) 10.1038/s41556-022-00882-3

[32] Matsuzaki, M., Honkura, N., Ellis-Davies, G.C.R., Kasai, H.: Structural basis of long-term potentiation in single dendritic spines. Nature 429(6993), 761–766 (2004)

[33] Okamoto, K.-I., Nagai, T., Miyawaki, A., Hayashi, Y.: Rapid and persistent modulation of actin dynamics regulates postsynaptic reorganization underlying bidirectional plasticity. Nat. Neurosci. 7(10), 1104–1112 (2004)

[34] Magnowska, M., Gorkiewicz, T., Suska, A., Wawrzyniak, M., Rutkowska-Wlodarczyk, I., Kaczmarek, L., Wlodarczyk, J.: Transient ECM protease activity promotes synaptic plasticity. Sci. Rep. 6(1), 27757 (2016)

[35] Cowan, A.E., Moraru, I.I., Schaff, J.C., Slepchenko, B.M., Loew, L.M.: Spatial modeling of cell signaling networks. Methods Cell Biol. 110, 195–221 (2012)

[36] Francis, E.A., Laughlin, J., Dokken, J.S., Finsberg, H., Lee, C.T., Rognes, M.E., Rangamani, P.: Spatial Modeling Algorithms for Reactions and Transport (SMART) in Biological Cells. bioRxiv (2024). 10.1101/2024.05.23.595604

[37] Novak, I.L., Gao, F., Choi, Y.-S., Resasco, D., Schaff, J.C., Slepchenko, B.M.: Diffusion on a curved surface coupled to diffusion in the volume: Application to cell biology. J. Comput. Phys. 226(2), 1271–1290 (2007)

[38] Lee, C.T., Laughlin, J.G., Angliviel de La Beaumelle, N., Amaro, R.E., McCammon, J.A., Ramamoorthi, R., Holst, M., Rangamani, P.: 3D mesh processing using GAMer 2 to enable reaction-diffusion simulations in realistic cellular geometries. PLoS Comput Biol 16(4), 1007756 (2020) 10.1371/journal.pcbi.1007756

[39] Helfrich, W.: Elastic Properties of Lipid Bilayers: Theory and Possible Experiments. Zeitschrift für Naturforschung C 28(11-12), 693–703 (1973) 10/gf3jnf

[40] Chen, Y., Saintillan, D., Rangamani, P.: Cell Motility Modes Are Selected by the Interplay of Mechanosensitive Adhesion and Membrane Tension. bioRxiv (2023). 10.1101/2023.05.31.543156

[41] Cao, Y., Ghabache, E., Rappel, W.-J.: Plasticity of cell migration resulting from mechanochemical coupling. Elife 8(e48478), 48478 (2019)

[42] Heinrich, L., Bennett, D., Ackerman, D., Park, W., Bogovic, J., Eckstein, N., Petruncio, A., Clements, J., Pang, S., Xu, C.S., Funke, J., Korff, W., Hess, H.F., Lippincott-Schwartz, J., Saalfeld, S., Weigel, A.V., COSEM Project Team: Whole-cell organelle segmentation in volume electron microscopy. Nature 599(7883), 141–146 (2021)

[43] Viana, M.P., Chen, J., Knijnenburg, T.A., Vasan, R., Yan, C., Arakaki, J.E., Bailey, M., Berry, B., Borensztejn, A., Brown, E.M., Carlson, S., Cass, J.A., Chaudhuri, B., Cordes Metzler, K.R., Coston, M.E., Crabtree, Z.J., Davidson, S., DeLizo, C.M., Dhaka, S., Dinh, S.Q., Do, T.P., Domingus, J., Donovan-Maiye, R.M., Ferrante, A.J., Foster, T.J., Frick, C.L., Fujioka, G., Fuqua, M.A., Gehring, J.L., Gerbin, K.A., Grancharova, T., Gregor, B.W., Harrylock, L.J., Haupt, A., Hendershott, M.C., Hookway, C., Horwitz, A.R., Hughes, H.C., Isaac, E.J., Johnson, G.R., Kim, B., Leonard, A.N., Leung, W.W., Lucas, J.J., Ludmann, S.A., Lyons, B.M., Malik, H., McGregor, R., Medrash, G.E., Meharry, S.L., Mitcham, K., Mueller, I.A., Murphy-Stevens, T.L., Nath, A., Nelson, A.M., Oluoch, S.A., Paleologu, L., Popiel, T.A., Riel-Mehan, M.M., Roberts, B., Schaefbauer, L.M., Schwarzl, M., Sherman, J., Slaton, S., Sluzewski, M.F., Smith, J.E., Sul, Y., Swain-Bowden, M.J., Tang, W.J., Thirstrup, D.J., Toloudis, D.M., Tucker, A.P., Valencia, V., Wiegraebe, W., Wijeratna, T., Yang, R., Zaunbrecher, R.J., Labitigan, R.L.D., Sanborn, A.L., Johnson, G.T., Gunawardane, R.N., Gaudreault, N., Theriot, J.A., Rafelski, S.M.: Integrated intracellular organization and its variations in human iPS cells. Nature 613(7943), 345–354 (2023)

[44] Hughes, T.J.R.: The Finite Element Method. Dover Civil and Mechanical Engineering. Dover Publications

[45] Ern, A., Guermond, J.-L.: Finite Elements III: First-Order and Time-Dependent PDEs. Texts in Applied Mathematics, vol. 74. Springer, Cham (2021). 10.1007/978-3-030-57348-5

[46] Belytschko, T., Liu, K., Moran, B., Elkhodary, K.: Nonlinear Finite Elements for Continua and Structures, 2nd edn. John Wiley & Sons, Nashville, TN (2013)

[47] Barrett, J.W., Garcke, H., Nürnberg, R.: Stable variational approximations of boundary value problems for Willmore flow with Gaussian curvature. IMA Journal of Numerical Analysis 37(4), 1657–1709 (2017) 10/gb4tvx

[48] Garcke, H., Nürnberg, R., Zhao, Q.: An energy-stable parametric finite element method for Willmore flow with normal-tangential velocity splitting. SIAM Journal on Scientific Computing 48(3), 1235–1259 (2026) 10.1137/25M1773878

[49] Mokbel, M., Mokbel, D., Liese, S., Weber, C., Aland, S.: A Simulation Method for the Wetting Dynamics of Liquid Droplets on Deformable Membranes. SIAM J. Sci. Comput. 46(6), 806–829 (2024) 10.1137/24M1641142

[50] Mercker, M., Marciniak-Czochra, A., Richter, T., Hartmann, D.: Modeling and Computing of Deformation Dynamics of Inhomogeneous Biological Surfaces. SIAM J. Appl. Math. 73(5), 1768–1792 (2013) 10.1137/120885553

[51] Ballatore, F., Madzvamuse, A., Jebane, C., Helfer, E., Allena, R.: A Geometric-Surface PDE Model for Cell-Nucleus Translocation through Confinement. bioRxiv (2025). 10.64898/2025.12.18.695144

[52] Dziuk, G., Elliott, C.M.: Finite element methods for surface PDEs. Acta Numerica 22, 289–396 (2013) 10/ggzzsq

[53] Mackenzie, J., Rowlatt, C., Insall, R.: A Conservative Finite Element ALE Scheme for Mass-Conservative Reaction-Diffusion Equations on Evolving Two-Dimensional Domains. SIAM J. Sci. Comput. 43(1), 132–166 (2021) 10/gh4xq8

[54] Happel, L., Voigt, A.: Toward a Two-Scale Model for Morphogenesis: How Cellular Processes Influence Tissue Deformations. Journal of Nonlinear Science 36(1), 15 (2026) 10.1007/s00332-025-10228-6

[55] Contri, A., Massing, A., Rangamani, P.: A finite element framework for solving coupled multiphysics problems with moving boundaries in cell biophysics. Computer Methods in Applied Mechanics and Engineering 459, 119071 (2026) 10.1016/j.cma.2026.119071

[56] Elliott, C.M., Ranner, T.: A unified theory for continuous-in-time evolving finite element space approximations to partial differential equations in evolving domains. IMA Journal of Numerical Analysis 41(3), 1696–1845 (2021) 10/gr8rfz

[57] Canham, P.B.: The minimum energy of bending as a possible explanation of the biconcave shape of the human red blood cell. Journal of Theoretical Biology 26(1), 61–81 (1970) 10/frc472

[58] Kovács, B., Power Guerra, C.A.: Higher order time discretizations with ALE finite elements for parabolic problems on evolving surfaces. IMA Journal of Numerical Analysis 38(1), 460–494 (2018) 10/gcz3xc

[59] Duan, B., Li, B.: New Artificial Tangential Motions for Parametric Finite Element Approximation of Surface Evolution. SIAM J. Sci. Comput. 46(1), 587–608 (2024) 10/gtj6hx

[60] Hansbo, P., Larson, M.G., Zahedi, S.: Characteristic cut finite element methods for convection–diffusion problems on time dependent surfaces. Computer Methods in Applied Mechanics and Engineering 293, 431–461 (2015) 10/f3pbxn

[61] Cheng, Q., Shen, J.: A New Lagrange Multiplier Approach for Constructing Structure Preserving Schemes, II. Bound Preserving. SIAM J. Numer. Anal. 60(3), 970–998 (2022) 10/gr5qjs

[62] Cheng, Q., Shen, J.: A new Lagrange multiplier approach for constructing structure preserving schemes, I. Positivity preserving. Computer Methods in Applied Mechanics and Engineering 391, 114585 (2022) 10/gr5qjr

[63] Hu, J., Li, B.: Evolving finite element methods with an artificial tangential velocity for mean curvature flow and Willmore flow. Numer. Math. 152(1), 127–181 (2022) 10/gr8rf4

[64] Labuz, E.C., Footer, M.J., Theriot, J.A.: Confined keratocytes mimic in vivo migration and reveal volume-speed relationship. Cytoskeleton (Hoboken) 80(1-2), 34–51 (2023)

[65] De Belly, H., Yan, S., Rocha, H., Ichbiah, S., Town, J.P., Zager, P.J., Estrada, D.C., Meyer, K., Turlier, H., Bustamante, C., Weiner, O.D.: Cell protrusions and contractions generate long-range membrane tension propagation. Cell 186(14), 3049–306115 (2023)

[66] Diz-Mu ñoz, A., Fletcher, D.A., Weiner, O.D.: Use the force: membrane tension as an organizer of cell shape and motility. Trends Cell Biol. 23(2), 47–53 (2013)

[67] Lieber, A.D., Yehudai-Resheff, S., Barnhart, E.L., Theriot, J.A., Keren, K.: Membrane tension in rapidly moving cells is determined by cytoskeletal forces. Curr. Biol. 23(15), 1409–1417 (2013)

[68] Francis, E.A., Heinrich, V.: Extension of chemotactic pseudopods by nonadherent human neutrophils does not require or cause calcium bursts. Sci. Signal. 11(521), 4289 (2018)

[69] Mankovich, A.R., Lee, C.-Y., Heinrich, V.: Differential effects of serum heat treatment on chemotaxis and phagocytosis by human neutrophils. PLoS One 8(1), 54735 (2013)

[70] Heinrich, V., Lee, C.-Y.: Blurred line between chemotactic chase and phagocytic consumption: an immunophysical single-cell perspective. J. Cell Sci. 124(Pt 18), 3041–3051 (2011)

[71] Lee, C.-Y., Thompson, G.R. 3rd, Hastey, C.J., Hodge, G.C., Lunetta, J.M., Pappagianis, D., Heinrich, V.: Coccidioides endospores and spherules draw strong chemotactic, adhesive, and phagocytic responses by individual human neutrophils. PLoS One 10(6), 0129522 (2015)

[72] Wangdi, T., Lee, C.-Y., Spees, A.M., Yu, C., Kingsbury, D.D., Winter, S.E., Hastey, C.J., Wilson, R.P., Heinrich, V., Bäumler, A.J.: The Vi capsular polysaccharide enables Salmonella enterica serovar typhi to evade microbe-guided neutrophil chemotaxis. PLoS Pathog. 10(8), 1004306 (2014)

[73] Zhelev, D.V., Alteraifi, A.M., Chodniewicz, D.: Controlled pseudopod extension of human neutrophils stimulated with different chemoattractants. Biophys. J. 87(1), 688–695 (2004)

[74] Herant, M., Marganski, W.A., Dembo, M.: The Mechanics of Neutrophils: Synthetic Modeling of Three Experiments. Biophys J 84(5), 3389–3413 (2003)

[75] Herant, M., Heinrich, V., Dembo, M.: Mechanics of neutrophil phagocytosis: behavior of the cortical tension. J. Cell Sci. 118(Pt 9), 1789–1797 (2005)

[76] Bell, M.K., Holst, M.V., Lee, C.T., Rangamani, P.: Dendritic spine morphology regulates calcium-dependent synaptic weight change. J. Gen. Physiol. 154(8), 202112980 (2022)

[77] Mesa, M.H., McCabe, K.J., Rangamani, P.: Synaptic cleft geometry modulates NMDAR opening probability by tuning neurotransmitter residence time. Biophys. J. 124(7), 1058–1072 (2025)

[78] Bourne, J.N., Harris, K.M.: Balancing structure and function at hippocampal dendritic spines. Annu. Rev. Neurosci. 31(1), 47–67 (2008)

[79] Holthoff, K., Tsay, D., Yuste, R.: Calcium dynamics of spines depend on their dendritic location. Neuron 33(3), 425–437 (2002)

[80] Wu, Y., Whiteus, C., Xu, C.S., Hayworth, K.J., Weinberg, R.J., Hess, H.F., De Camilli, P.: Contacts between the endoplasmic reticulum and other membranes in neurons. Proc. Natl. Acad. Sci. U. S. A. 114(24), 4859–4867 (2017)

[81] Bonilla-Quintana, M., Rangamani, P.: Biophysical Modeling of Actin-Mediated Structural Plasticity Reveals Mechanical Adaptation in Dendritic Spines. eNeuro 11(3) (2024) 10.1523/ENEURO.0497-23.2024 . Chap. Research Article: New Research

[82] Rangamani, P., Levy, M.G., Khan, S., Oster, G.: Paradoxical signaling regulates structural plasticity in dendritic spines. Proc. Natl. Acad. Sci. U. S. A. 113(36), 5298–307 (2016)

[83] Bell, M., Bartol, T., Sejnowski, T., Rangamani, P.: Dendritic spine geometry and spine apparatus organization govern the spatiotemporal dynamics of calcium. J. Gen. Physiol. 151(8), 1017–1034 (2019)

[84] Crane, K., Weischedel, C., Wardetzky, M.: The heat method for distance computation. Commun. ACM 60(11), 90–99 (2017) 10/gcj3hk

[85] Bosch, M., Castro, J., Saneyoshi, T., Matsuno, H., Sur, M., Hayashi, Y.: Structural and molecular remodeling of dendritic spine substructures during long-term potentiation. Neuron 82(2), 444–459 (2014)

[86] Johnson, M.E., Chen, A., Faeder, J.R., Henning, P., Moraru, I.I., Meier-Schellersheim, M., Murphy, R.F., Prüstel, T., Theriot, J.A., Uhrmacher, A.M.: Quantifying the roles of space and stochasticity in computer simulations for cell biology and cellular biochemistry. Mol. Biol. Cell 32(2), 186–210 (2021)

[87] Xiong, Y., Rangamani, P., Fardin, M.-A., Lipshtat, A., Dubin-Thaler, B., Rossier, O., Sheetz, M.P., Iyengar, R.: Mechanisms controlling cell size and shape during isotropic cell spreading. Biophys J 98(10), 2136–2146 (2010) 10.1016/j.bpj.2010.01.059

[88] Fardin, M.A., Rossier, O.M., Rangamani, P., Avigan, P.D., Gauthier, N.C., Vonnegut, W., Mathur, A., Hone, J., Iyengar, R., Sheetz, M.P.: Cell spreading as a hydrodynamic process. Soft Matter 6, 4788–4799 (2010)

[89] MacDonald, G., Mackenzie, J.A., Nolan, M., Insall, R.H.: A computational method for the coupled solution of reaction-diffusion equations on evolving domains and manifolds. J. Comput. Phys. 309(C), 207–226 (2016) 10.1016/j.jcp.2015.12.038

[90] Bachini, E., Krause, V., Nitschke, I., Voigt, A.: Derivation and simulation of a two-phase fluid deformable surface model. Journal of Fluid Mechanics 977, 41 (2023) 10.1017/jfm.2023.943

[91] Ethridge, F., Greengard, L.: A new fast-multipole accelerated Poisson solver in two dimensions. SIAM J. Sci. Comput. 23(3), 741–760 (2001)

[92] Höllein, T., Aland, S.: A phase-field model to simulate membrane remodeling and topology changes induced by wetting droplets. Comput. Methods Appl. Mech. Eng. 456(118938), 118938 (2026)

[93] Liang, Y., Celiker, E., Lin, P.: A phase-field model for vesicle membranes incorporating area-difference elasticity. PLoS Comput. Biol. 22(4), 1014185 (2026)

[94] Zhang, T., Wolgemuth, C.W.: A general computational framework for the dynamics of single- and multi-phase vesicles and membranes. J. Comput. Phys. 450(110815), 110815 (2022)

[95] Bottacchiari, M., Gallo, M., Bussoletti, M., Casciola, C.M.: The local variation of the gaussian modulus enables different pathways for fluid lipid vesicle fusion. Sci. Rep. 14(1), 23 (2024)

[96] Burman, E., Claus, S., Hansbo, P., Larson, M.G., Massing, A.: CutFEM: Discretizing geometry and partial differential equations. International Journal for Numerical Methods in Engineering 104(7), 472–501 (2015) 10.1002/nme.4823

[97] Ulfsby, T.B., Massing, A., Sticko, S.: Stabilized cut discontinuous Galerkin methods for advection–reaction problems on surfaces. Computer Methods in Applied Mechanics and Engineering 413, 116109 (2023) 10/gsb7hg

[98] Reiner, J., Linden, N., Vaziri, R., Zobeiry, N., Kramer, B.: Bayesian parameter estimation for the inclusion of uncertainty in progressive damage simulation of composites. Compos. Struct. 321(117257), 117257 (2023)

[99] Xun, X., Cao, J., Mallick, B., Carroll, R.J., Maity, A.: Parameter estimation of partial differential equation models. J. Am. Stat. Assoc. 108(503), 1009–1020 (2013)

[100] Vanslambrouck, M., Thiels, W., Vangheel, J., Bavel, C., Smeets, B., Jelier, R.: Image-based force inference by biomechanical simulation. PLoS Comput. Biol. 20(12), 1012629 (2024)

[101] Jurado, A., Isensee, J., Hofemeier, A., Krüger, L.J., Wittkowski, R., Golestanian, R., Bittihn, P., Betz, T.: 3D multiscale shape analysis of nuclei and in vivo elastic stress sensors allows force inference. Biophys. J. 124(17), 2784–2796 (2025)

[102] Barrett, J.W., Garcke, H., Nürnberg, R.: Chapter 4 - Parametric finite element approximations of curvature-driven interface evolutions. In: Bonito, A., Nochetto, R.H. (eds.) Handbook of Numerical Analysis. Geometric Partial Differential Equations -Part I, vol. 21, pp. 275–423. Elsevier, Amsterdam (2020). 10.1016/bs.hna.2019.05.002

[103] Alphonse, A., Elliott, C.M., Stinner, B.: An abstract framework for parabolic PDEs on evolving spaces. Portugaliae Mathematica 72(1), 1–46 (2015) 10.4171/pm/1955

[104] Alphonse, A., Caetano, D., Djurdjevac, A., Elliott, C.M.: Function spaces, time derivatives and compactness for evolving families of Banach spaces with applications to PDEs. Journal of Differential Equations 353, 268–338 (2023) 10.1016/j.jde.2022.12.032

[105] Ern, A., Guermond, J.-L.: Finite Elements II: Galerkin Approximation, Elliptic And Mixed PDEs. Texts in Applied Mathematics, vol. 73. Springer, Cham (2021). 10.1007/978-3-030-56923-5

[106] Hirt, C.W., Amsden, A.A., Cook, J.L.: An arbitrary Lagrangian-Eulerian computing method for all flow speeds. Journal of Computational Physics 14(3), 227–253 (1974) 10/fdtb8s

[107] Hughes, T.J.R., Liu, W.K., Zimmermann, T.K.: Lagrangian-Eulerian finite element formulation for incompressible viscous flows. Computer Methods in Applied Mechanics and Engineering 29(3), 329–349 (1981) 10/fjv3wd

[108] Nobile, F.: Numerical approximation of fluid-structure interaction problems with application to haemodynamics. PhD thesis, EPFL (2001). 10.5075/epfl-thesis-2458

[109] Gastaldi, L.: A priori error estimates for the Arbitrary Lagrangian Eulerian formulation with finite elements 9(2), 123–156 (2001) 10.1515/JNMA.2001.123. Chap. Journal of Numerical Mathematics

[110] Formaggia, L., Nobile, F.: Stability analysis of second-order time accurate schemes for ALE–FEM. Computer Methods in Applied Mechanics and Engineering 193(39), 4097–4116 (2004) 10/cvf4fz

[111] Badia, S., Codina, R.: Analysis of a stabilized finite element approximation of the transient convection-diffusion equation using an ALE framework (2006)

[112] Bonito, A., Kyza, I., Nochetto, R.H.: Time-discrete higher order ALE formulations: A priori error analysis. Numer. Math. 125(2), 225–257 (2013) 10/f49ctb

[113] Bänsch, E., Deckelnick, K., Garcke, H., Pozzi, P.: Interfaces: Modeling, Analysis, Numerics. Oberwolfach Seminars, vol. 51. Springer, Cham (2023). 10.1007/978-3-031-35550-9

[114] Barrett, J.W., Garcke, H., Nürnberg, R.: A parametric finite element method for fourth order geometric evolution equations. Journal of Computational Physics 222(1), 441–467 (2007) 10/cr3n7j

[115] Boffi, D., Gastaldi, L.: Stability and geometric conservation laws for ALE formulations. Computer Methods in Applied Mechanics and Engineering 193(42), 4717–4739 (2004) 10.1016/j.cma.2004.02.020

[116] Nobile, F., Formaggia, L.: A Stability Analysis for the Arbitrary Lagrangian Eulerian Formulation with Finite Elements 7(2), 105–132 (1999)

[117] Barreira, R., Elliott, C.M., Madzvamuse, A.: The surface finite element method for pattern formation on evolving biological surfaces. J. Math. Biol. 63(6), 1095–1119 (2011) 10/b386pd

[118] Kovács, B., Li, B., Lubich, C., Power Guerra, C.A.: Convergence of finite elements on an evolving surface driven by diffusion on the surface. Numer. Math. 137(3), 643–689 (2017) 10/gh4xvx

[119] Zhu, C., Lee, C.T., Rangamani, P.: Mem3DG: Modeling membrane mechanochemical dynamics in 3D using discrete differential geometry. Biophysical Reports 2(3), 100062 (2022) 10/gsfcr2

[120] Schöberl, J.: C++11 Implementation of Finite Elements in NGSolve, 1–23 (2014)

[121] Pollard, T.D., Blanchoin, L., Mullins, R.D.: Molecular mechanisms controlling actin filament dynamics in nonmuscle cells. Annu. Rev. Biophys. Biomol. Struct. 29(1), 545–576 (2000)

[122] Sase, I., Miyata, H., Ishiwata, S., Kinosita, K. Jr: Axial rotation of sliding actin filaments revealed by single-fluorophore imaging. Proc. Natl. Acad. Sci. U. S. A. 94(11), 5646–5650 (1997)

[123] Rock, R.S., Rief, M., Mehta, A.D., Spudich, J.A.: In vitro assays of processive myosin motors. Methods 22(4), 373–381 (2000)

[124] McGrath, J.L., Tardy, Y., Dewey, C.F. Jr, Meister, J.J., Hartwig, J.H.: Simultaneous measurements of actin filament turnover, filament fraction, and monomer diffusion in endothelial cells. Biophys. J. 75(4), 2070–2078 (1998)

[125] Herant, M., Heinrich, V., Dembo, M.: Mechanics of neutrophil phagocytosis: Experiments and quantitative models. Journal of Cell Science 119(9), 1903–1913 (2006) 10/bzxf63

[126] Zhelev, D.V., Needham, D., Hochmuth, R.M.: Role of the membrane cortex in neutrophil deformation in small pipets. Biophys. J. 67(2), 696–705 (1994)

